# Cell-Type Specific Reductions in Interneuron Gene Expression within Subregions of the Anterior and Posterior Cingulate Gyrus of Schizophrenia and Bipolar Disorder Subjects

**DOI:** 10.1101/2024.03.20.585908

**Authors:** David M. Krolewski, Maria Waselus, Blynn G. Bunney, Richard M. Myers, Jack D. Barchas, Francis S.Y. Lee, Alan F. Schatzberg, William E. Bunney, Huda Akil, Stanley J. Watson

## Abstract

Schizophrenia (SZ) and bipolar disorder (BP) patients share overlapping and distinct neurocognitive deficits. Neuroimaging of these patients and postmortem gene expression analyses suggest that compromised cingulate gyrus GABA-ergic interneurons may contribute to cognitive impairments in SZ and BP. Therefore, we investigated potential gene expression signatures for SZ and BP using interneuron cell-type specific markers including glutamic acid decarboxylase (GAD67), parvalbumin (PV), somatostatin (SST), and vasoactive intestinal peptide (VIP) within cingulate Brodmann’s areas (BA). We report reduced GAD67 mRNA in anterior midcingulate cortex (aMCC) of SZ and BP subjects with BA24c’ being most dysregulated across disorders, demonstrating reduced PV (SZ), SST (BP), and VIP mRNA (SZ and BP). Dorsal posterior cingulate (dPCC) displayed decreased SST (BP) whereas retrosplenial cortex (RSC) showed reduced PV (SZ and BP) and SST mRNA (BP). Our results show shared and unique transcription signatures of two disorders in specific cingulate gyrus regions and cell types. SZ and BP show a similar profile of aMCC gene expression reductions suggesting subregional dysregulation within areas associated with error/action monitoring and the saliency network. In dPCC/RSC, transcriptional changes are associated with disease-specific gene/subregion signatures, possibly underlying differential effects on allocation of attentional resources and visuospatial memory processing unique to each disorder.

## Introduction

The cingulate gyrus is a functionally heterogeneous cortical structure that mediates various forms of neurocognition including motor control, attention, and memory processing (1,2) which are reported to be compromised in schizophrenia (SZ) and bipolar disorder (BP) (3,4). Coordination of cognitive domains is achieved via cytoarchitecturally parcellated subdivisions, or Brodmann’s areas (BA) (5). These include the anterior midcingulate cortex (aMCC; BA32’, 24’), dorsal posterior cingulate cortex (dPCC; BAd31, d23), and retrosplenial cortex (RSC; BA30, 29). Often described as dorsal ACC in literature, aMCC mediates error processing, action monitoring, and evaluative learning from action-outcomes concerning reward engagement and pain/fear avoidance behaviors (6,7). dPCC is linked to recognition memory processing and attention set shifting (8,9). RSC is associated with spatial navigation and memory retrieval (10,11).

Despite being highly interconnected (1,2), imaging studies demonstrate integration of aMCC, dPCC, and RSC modular operations through distinct hubs within different resting-state large-scale neurocognitive networks (12,13). dPCC/RSC are components of the default mode network (DMN) characterized by its heightened metabolic status during internally directed attention in the absence of stimulus-driven tasks (14–16). DMN activation drives self-referential constructs including mentalizing, future planning, and autobiographic/episodic memory (17,18). The aMCC-anchored saliency network (SAN) directs attention toward pertinent stimuli, attenuating DMN activity and thereby limiting self-referentiality in favor of filtering interoceptive-autonomic, emotion, and reward processing information (12,19). Higher order goal-directed tasks, e.g., involving working memory, recruit the dorsal lateral prefrontal cortex (DLPFC)-driven central executive network (CEN) which has capacity to bridge with the SAN, co-acting to maintain physiological and stimulus saliency (12,19). According to the triple network model of psychopathology, task information processing is facilitated by SN-mediated switching between the DMN and CEN based upon cognitive demand (12) whereas aberrant inter/intra-network functional connectivity (FC) is associated with cognitive impairments in SZ and BP patients (20,21).

Evidence suggests that dysfunction of aMCC inhibitory gamma aminobutyric acid (GABA)-ergic interneurons may disrupt local and large-scale networks potentially lending to the neurocognitive manifestations of psychiatric illness. Meta-analyses show that patients with SZ consistently display decreased aMCC GABA levels (22,23). Functional imaging studies demonstrate abnormal associations between aMCC GABA and, as part of the SAN, FC with DMN/CEN nodes in SZ (24–26). Elucidating the pathological underpinnings of proposed dysregulation of cingulate GABA-ergic neurocircuits in SZ has been aided by examination of discrete interneuron subpopulations. Three largely separate GABA-ergic cell-types concordantly producing the GABA synthesizing enzyme glutamic acid decarboxylase (GAD67) with either the calcium-binding protein parvalbumin (PV), neuropeptide somatostatin (SST), or vasoactive intestinal peptide (VIP), exhibit differential synaptic connectivity and collectively coordinate excitatory pyramidal neuron activity (27). As a result, compromised function of any single specific cell-type may reshape corticocortical communication. Indeed, aligned with an aberrant SAN, postmortem gene expression analyses in aMCC of SZ subjects reveal diminished GAD67, PV, and SST mRNA (28,29), suggesting that alteration of both local inhibitory neurocircuits and corresponding pyramidal cell output may contribute to associated neurocognitive deficits.

Notwithstanding compelling evidence connecting aMCC GABA-ergic interneurons to the psychopathology of SZ, there is a clear need for further assessment of the posterior cingulate gyrus in SZ. There is also a dearth of evidence around the dysregulation in BP overall. Thus, we conducted the present in situ hybridization study to evaluate GAD67, PV, SST, and VIP gene expression within distinct BAs of the anterior and posterior cingulate gyrus in SZ and BP postmortem subjects. The results shed further light on the potential mechanisms underlying features of neurocognitive impairment in two psychiatric disorders.

## Materials and Methods

### Sample collection

Postmortem brain tissue from a total of 72 subjects diagnosed as normal CTR, SZ, or BP were obtained from the Pritzker Brain Bank at the University of California, Irvine (UCI). All studies were approved by the Institutional Review Board of UCI and the University of Michigan. All subjects had zero agonal factors with postmortem interval of <37 hours and pH above 6.5 to ensure mRNA preservation (30,31) (see summaries **Table 1**, also comprehensive demographics **Tables S1 and S2**).

**Table 1:**
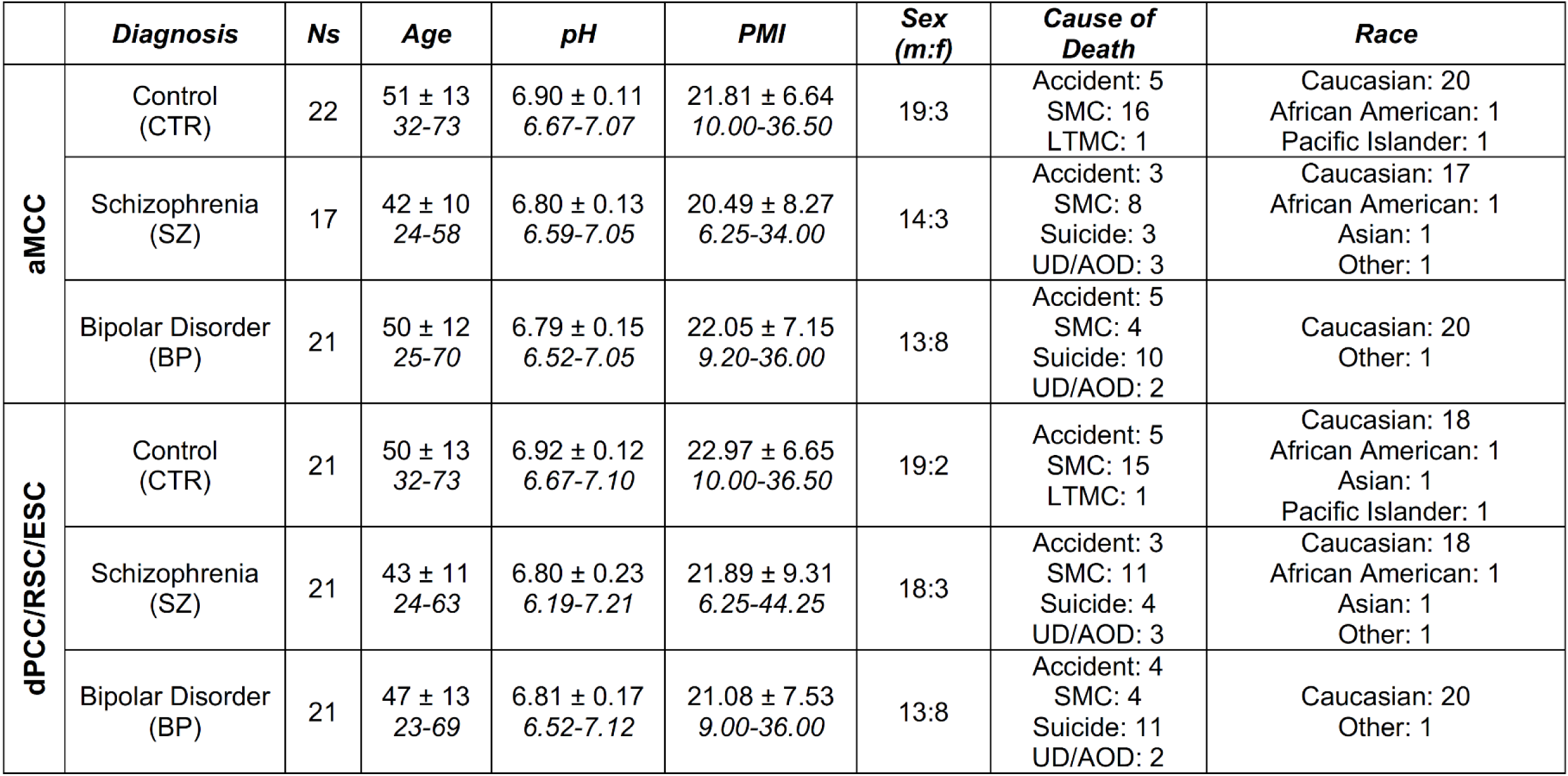
Demographic information for diagnostic groups included in the aMCC and dPCC/RSC/ESC analyses. Summary values are listed for age, pH, and PMI (mean ± standard deviation) for the aMCC and dPCC/RSC/ESC regions by diagnosis, with the range of values for each group italicized below (when appropriate; also, see **Tables S1** and **S2** and **Fig. S2**). aMCC: anterior midcingulate cortex; dPCC: dorsal posterior cingulate cortex; RSC: retrosplenial cortex; ESC: ectosplenial cortex; n: number; m: male; f: female; SMC: sudden medical condition; UD/AOD: undetermined/accidental overdose; LTMC: long-term medical condition; PMI: postmortem interval

|  | *Diagnosis* | *Ns* | *Age* | *pH* | *PMI* | *Sex (m:f)* | *Cause of Death* | *Race* |
| --- | --- | --- | --- | --- | --- | --- | --- | --- |
| **aMCC** | Control (CTR) | 22 | 51 ± 13<br>32-73 | 6.90 ± 0.11<br>6.67-7.07 | 21.81 ± 6.64<br>10.00-36.50 | 19:3 | Accident: 5<br>SMC: 16<br>LTMC: 1 | Caucasian: 20<br>African American: 1<br>Pacific Islander: 1 |
|  | Schizophrenia (SZ) | 17 | 42 ± 10<br>24-58 | 6.80 ± 0.13<br>6.59-7.05 | 20.49 ± 8.27<br>6.25-34.00 | 14:3 | Accident: 3<br>SMC: 8<br>Suicide: 3<br>UD/AOD: 3 | Caucasian: 17<br>African American: 1<br>Asian: 1<br>Other: 1 |
|  | Bipolar Disorder (BP) | 21 | 50 ± 12<br>25-70 | 6.79 ± 0.15<br>6.52-7.05 | 22.05 ± 7.15<br>9.20-36.00 | 13:8 | Accident: 5<br>SMC: 4<br>Suicide: 10<br>UD/AOD: 2 | Caucasian: 20<br>Other: 1 |
| **dPCC/RSC/ESC** | Control (CTR) | 21 | 50 ± 13<br>32-73 | 6.92 ± 0.12<br>6.67-7.10 | 22.97 ± 6.65<br>10.00-36.50 | 19:2 | Accident: 5<br>SMC: 15<br>LTMC: 1 | Caucasian: 18<br>African American: 1<br>Asian: 1<br>Pacific Islander: 1 |
|  | Schizophrenia (SZ) | 21 | 43 ± 11<br>24-63 | 6.80 ± 0.23<br>6.19-7.21 | 21.89 ± 9.31<br>6.25-44.25 | 18:3 | Accident: 3<br>SMC: 11<br>Suicide: 4<br>UD/AOD: 3 | Caucasian: 18<br>African American: 1<br>Asian: 1<br>Other: 1 |
|  | Bipolar Disorder (BP) | 21 | 47 ± 13<br>23-69 | 6.81 ± 0.17<br>6.52-7.12 | 21.08 ± 7.53<br>9.00-36.00 | 13:8 | Accident: 4<br>SMC: 4<br>Suicide: 11<br>UD/AOD: 2 | Caucasian: 20<br>Other: 1 |

### Brain dissection and in situ hybridization (ISH)

Slabs containing the aMCC and dPCC/RSC/ectosplenial (ESC) regions were visually identified with respect to gross anatomical landmarks and subdissected (see Supplemental Methods and **Fig. S1** for representative sagittal brain schematic illustrating gross cingulate dissection locations). Tissue processing methods have been previously described (32). Briefly, following brain removal, coronal slabs of ∼0.8 cm were cut, flash-frozen, and stored at -80°C. Blocks were sectioned at 10 µm in a cryostat, thaw mounted onto Superfrost^TM^ Plus glass slides (Fisher Scientific, Pittsburgh, PA), and stored at -80°C. ^35^S-labeled ISH was performed utilizing gene-specific [^35^S]UTP and [^35^S]ATP-labeled cRNA riboprobes for GAD67, PV, SST, and VIP. Slides containing the sections were exposed to film and developed at which point digitized images were quantified with ImageJ software (NIH) to measure signal intensity for each gene. Additional details regarding ISH, quantitative assessment, and image preparation/manipulation are included in Supplemental Methods.

### Statistical analysis

Statistical analyses were completed using IBM SPSS Statistics (Version 27) and graphs prepared using GraphPad Prism version 9.3.1 for Windows (GraphPad Software, San Diego, California USA, www.graphpad.com). Demographic variables of age, pH, and PMI were analyzed using a one-way ANOVA. Gene expression in control subjects was examined using a two-way ANOVA where anatomical subregion and gene were included as factors. A linear mixed effects model was used to evaluate whether diagnosis of SZ or BP resulted in gene expression changes in aMCC or dPCC/RSC/ESC subregions. The dependent variable, o.d., was stratified by the fixed effects of diagnosis, cingulate subregion, and gene. Subject was included as a random effect and pH as a covariate. For all described analyses, Bonferroni post-hoc correction was used when appropriate. All results were considered statistically significant at p<0.05.

## Results

### Demographic variables

Subject demographics included age, pH, and PMI. Comparisons between CTR, SZ, and BP subject groups in the aMCC and dPCC/RSC/ESC (see **Table 1**, also **Tables S1** and **S2**) indicated that brain pH was significantly different between groups in the aMCC (**Fig. S2B**; F(2,57)=4.804, p=0.012), with lower brain pH found in BP (pH=6.79) vs. CTR (pH=6.9) subjects (BP < CTR; *p*=0.020). Consequently, pH was included as a covariant in subsequent statistical analyses.

### Gene Expression in Control Subjects

Visual inspection of ISH signal demonstrated anticipated distributions for each gene, along with PV gradients in the aMCC and dPCC/RSC/ESC regions of control CTR subjects **(Figs. 1,2)**. The distribution of PV appeared to be more intense in the dorsal aspects of the aMCC (**Fig. 1C,D**) and stronger ventrally within the posterior regions (**Fig. 2C,E**), particularly within the RSC/ESC. In the aMCC, these observations were supported by a significant gene x subregion interaction in CTR (F(6,208) = 2.965, p=0.008) which indicated that PV expression differed by aMCC subregion (F(2,208) = 8.019, p<0.001). PV expression was lower in BA24ab’ compared to BA32’ (**Fig. 1G**; p<0.001) and BA24c’ (**Fig. 1G**; p=0.018). Similarly, the interaction between gene and subregion in the dPCC/RSC/ESC of CTR subjects was significant (F(15,366) =15,366) =20.640, p<0.001), and only PV expression varied across dPCC/RSC/ESC subregions (F(5,366) = 74.664, p<0.001). In the dPCC/RSC/ESC, there was dorsoventral and mediolateral gradient of PV expression, with the lowest PV levels measured dorsally in BAd23c and BAd23b **(Fig. 2C,E,F, Table S3)**. Ventrally, PV intensity in BAd23a was higher vs. BAd23c and BAd23b (**Fig. 2C,E,F, Table S3;** p<0.05). RSC BA30 demonstrated elevated PV vs. BAd23c, BAd23b, and BAd23a (**Fig. 2C,E,F, Table S3;** p<0.001). Lastly, BA29 and BA26 were similar (p>0.05), expressing the highest PV o.d. compared to all other regions (**Fig. 2C,E,F, Table S3;** p<0.001). No variations in GAD67, SST, or VIP o.d. were found between subregions in the aMCC (**Fig. S3 A,C,E**) or dPCC/RSC/ESC of CTRs (**Fig. S3 B,D,F**).

**Figure 1.**
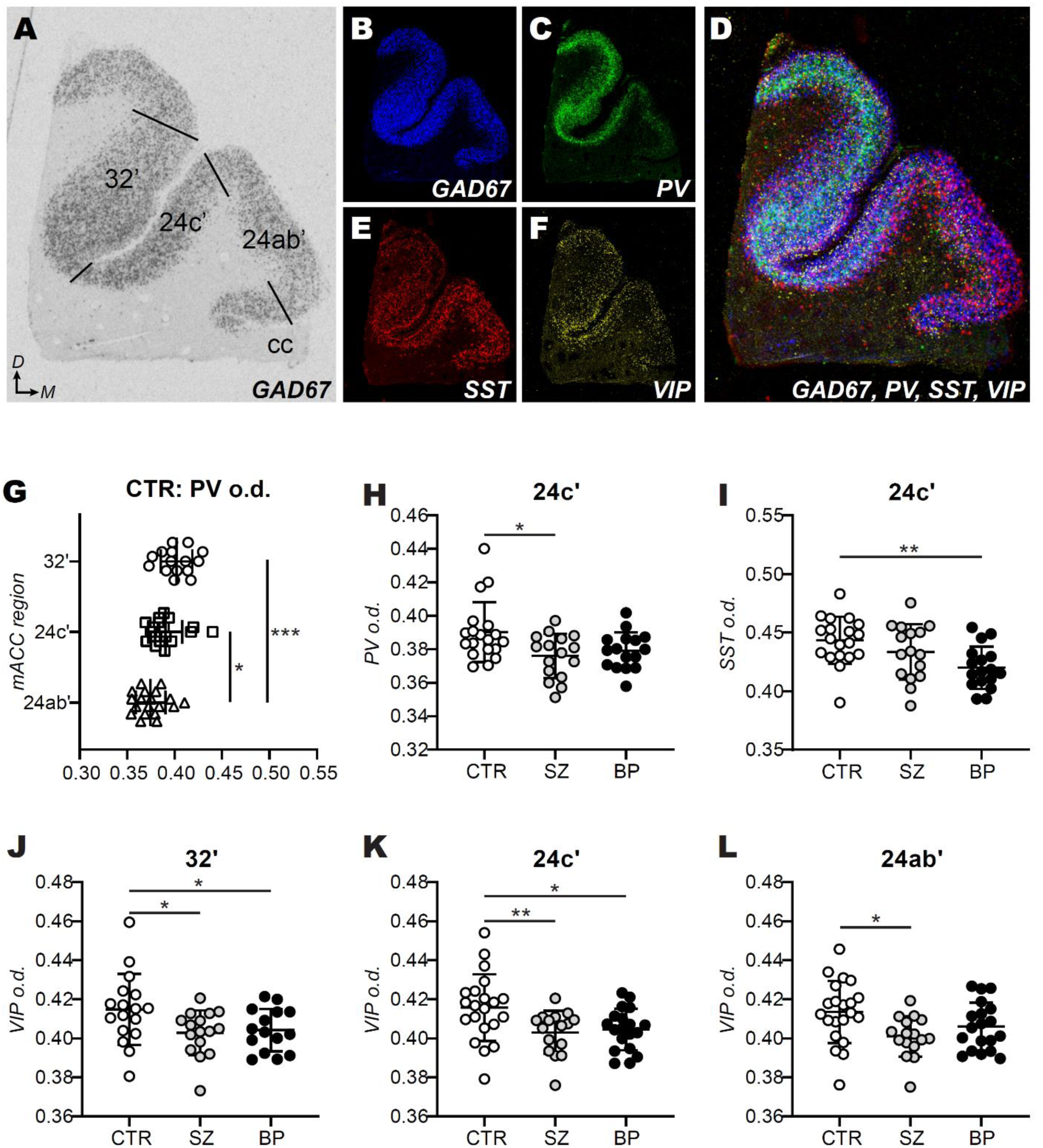
Interneuron markers exhibit different distribution patterns in aMCC subregions as well as differences in gene expression between healthy control vs. psychiatric groups as revealed by radioactive in situ hybridization. **A)** GAD67 mRNA expression is shown in regions 32’, 24c’ and 24ab’ of the aMCC. **B-E)** Monocolor images of adjacent in situ hybridization sections were pseudocolored, aligned and **F)** overlayed to examine the relative spatial distributions in gene expression as shown. Note that the distribution of PV within aMCC tissue sections was most varied across subregions (**C,D**). **G)** In healthy control subjects, the average aMCC PV mRNA expression was reduced in region 24ab’ compared to region 32’ (p<0.001) and 24c’ (p=0.018). **H)** SZ subjects had reduced levels of PV vs. CTR, specifically in region 24c’ (p=0.022). **I)** Reduced SST expression was also noted in 24c’, where BP subjects had reduced expression vs CTR (p=0.003). **J-L)** VIP expression was lower in SZ subjects vs CTR in all examined aMCC subregions (**J**: p=0.024; **K**: p=0.010; **L**: p=0.015), while BP subjects also had reduced VIP mRNA in regions 32’ (**J**: p=0.011) and 24c’ (**K**: p=0.026) vs. CTR. Each dot (**G-L**) represents one subject. Data is presented as mean +/-standard deviation. ***p<0.001; **p<0.01; *p<0.05 aMCC: anterior midcingulate cortex; cc: corpus callosum; *D*: dorsal; GAD67: glutamic acid decarboxylase 67; *M*: medial; PV: parvalbumin; SST: somatostatin; VIP: vasoactive intestinal peptide

**Figure 2.**
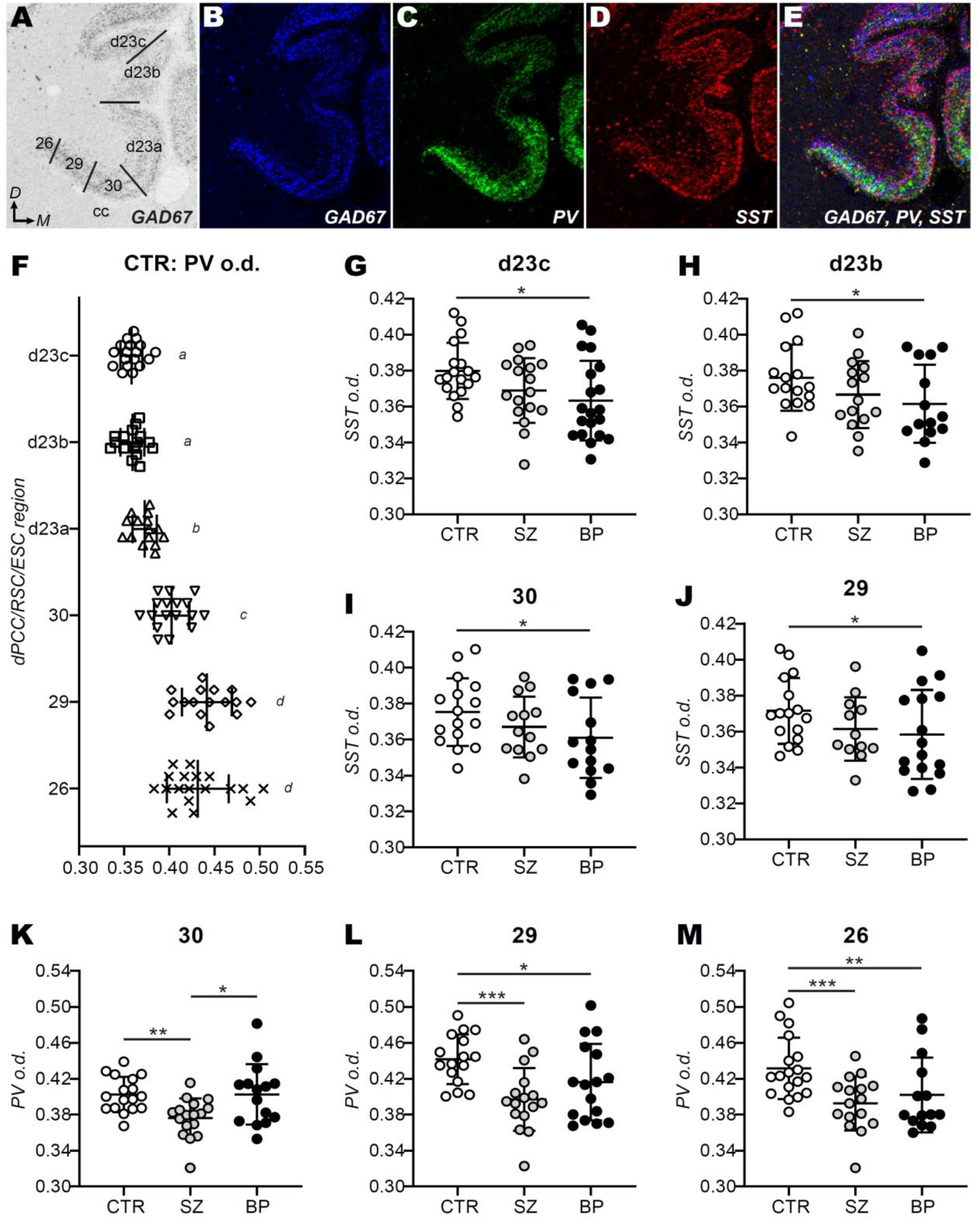
Distribution and differential expression of interneuron marker genes within the dPCC/RSC/ESC of psychiatric and control subjects. **A)** GAD67 mRNA expression is shown in the dPCC (regions d23c, d23b, and d23a), the RSC (regions 30 and 29) and ESC (region 26). **B-D)** Monochrome images of adjacent tissues sections processed for in situ hybridization were pseudocolored, aligned and **E)** overlayed using Adobe Photoshop to visually examine the relative distribution of signal (VIP not shown). **F)** There was a gradient of PV expression in the dPCC/RSC/ESC, with regions d23c and d23b having the lowest (and comparable, “*a*”) levels of PV expression and regions 29 and 26 exhibiting the most robust (and comparable, “*d*”) expression (**C,E**). Significant differences in PV expression between dPCC/RSC/ESC subregions are indicated by different italicized letters (see **Table S3** for detailed group differences). **G-H)** In the dPCC, only SST expression differed significantly between diagnoses, where BP subjects exhibited reduced SST vs. CTRs in region d23c (**G**; p=0.014) and d23b (**H**; p=0.013). **I-J)** SST expression was also significantly reduced in BP vs. control subjects in the RSC (**I**: p=0.014; **J**: p=0.016). **K-M)** PV mRNA differed between diagnoses in the RSC and ESC, where SZ subjects had decreased PV in both the RSC (**K**: SZ vs. CTR: p=0.005; SZ vs. BP: p=0.034; and **L**: SZ vs. CTR: p<0.001) and ESC (**M**: p=0.001). PV expression was also decreased in BP subjects compared to CTRs in RSC region 29 (**L**: p=0.013) and ESC (**M**: p=0.009). Each dot represents one subject; data is presented as mean +/-standard deviation. ***p<0.001; **p<0.01; *p<0.05 BP: Bipolar Disorder subject group; cc: corpus callosum; CTR: non-psychiatric control subject group; *D*: dorsal; dPCC: dorsal posterior cingulate cortex; ESC: ectosplenial cortex; GAD67: glutamic acid decarboxylase 67; *M*: medial; PV: parvalbumin; RSC: retrosplenial cortex; SST: somatostatin; SZ: schizophrenia subject group

**Figure 3.**
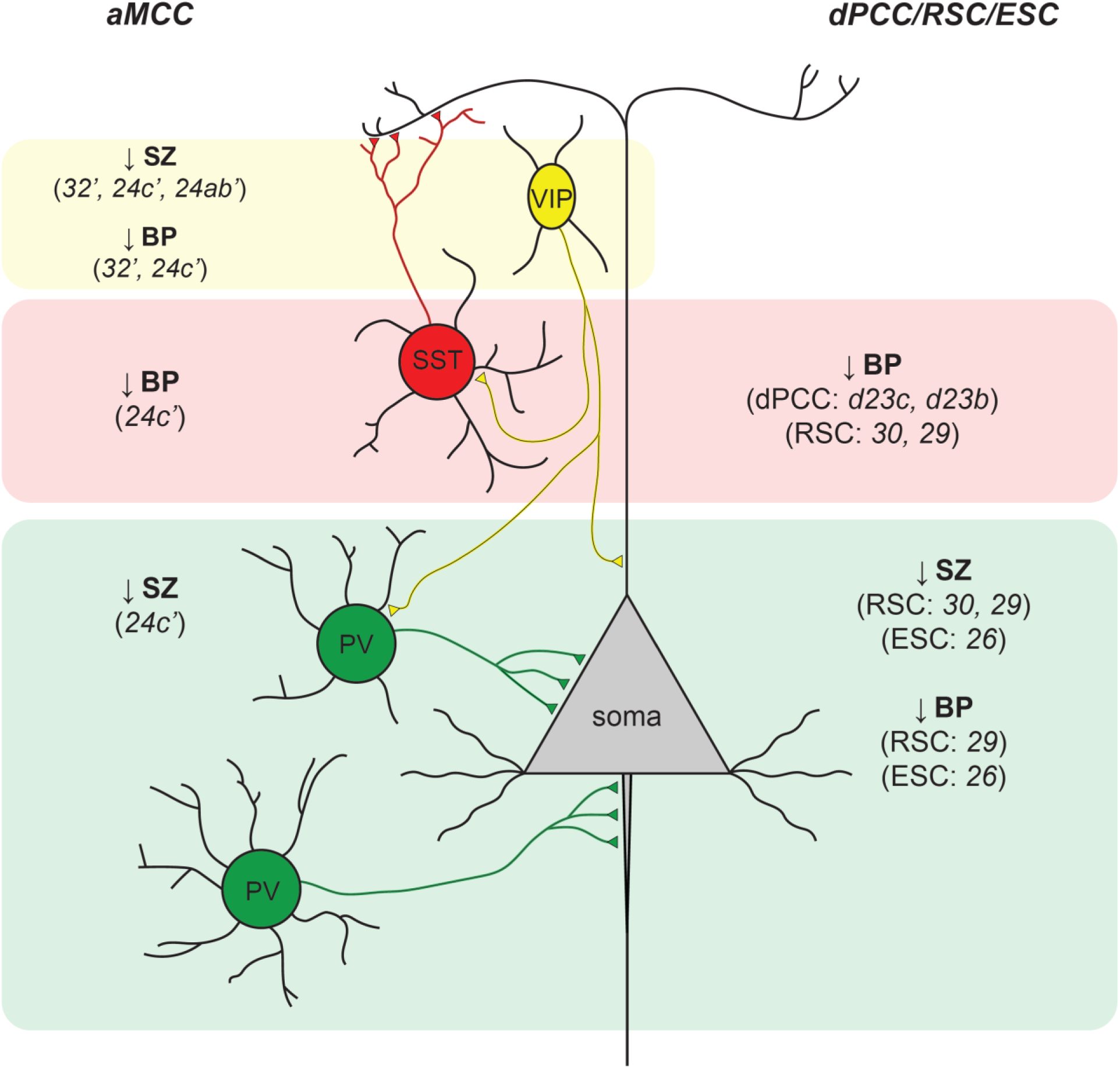
Summary of interneuron-related gene expression changes. Decreases in the expression of interneuron-related genes (top to bottom, VIP: yellow, SST: red; PV: green) are summarized for both the aMCC (left) and dPCC/RSC/ESC (right) cortical regions. In the aMCC, VIP expression was decreased in both SZ (32’, 24c’, and 24ab’) and BP (regions 32’ and 24c’) subjects compared to CTR, presumably reflecting a disinhibition of cortical interneurons (e.g., SST and/or PV) as well as decreased inhibition at the apical dendrite of pyramidal neurons. Compared to CTRs, SST expression was decreased in BP subjects in the aMCC (24c’) and both the dPCC (regions d23c and d23b) and RSC (regions 30 and 29). These decreases in SST likely occur in Martinotti cells, known to synapse onto the apical tufts of excitatory pyramidal neurons. Decreased PV expression (vs. CTRs) was found in SZ subjects in the aMCC (24c’), RSC (regions 30 and 29) and ESC (region 26) while decreased PV expression in BP subjects (vs CTRs) was only found in the RSC region 29 and ESC (region 26). PV expression is localized to both basket cells and chandelier cells of the aMCC and RSC/ESC where these neurons contact pyramidal neurons at the soma and initial axon segment, respectively.

### Diagnosis-specific aMCC gene expression

We first determined if aMCC gene expression was different between the CTR, SZ and BP groups. The effect of diagnosis was significant (F(2,79.312) = 5.872, p=0.004) and indicated interneuron o.d. measurements were lower in SZ (p=0.020) and BP (p=0.009) groups versus CTR (data not shown). To determine whether the effect of diagnosis was specific to a particular gene, we looked at the interaction between diagnosis and gene and determined that it was significant (F(9, 169.658) = 92.920, p<0.001). GAD67 (F(2, 171.706) = 4.307, p=0.015), SST (F(2,166.427) = 5.272, p=0.006), and VIP (F(2,58.233) = 3.994, p=0.024) expression all differed by diagnosis, with both SZ (GAD67: p=0.020; SST: p=0.002, VIP: p=0.038) and BP (GAD67: p=0.009; SST: p=0.043; VIP: p=0.010) subjects exhibiting lower mRNA expression than CTR. Differences in PV expression between CTR, SZ and BP groups was not significant (F(2,124.406) = 3.000, p=0.053).

When examining potential differences in specific genes and their subregional restriction within the aMCC, a significant 3-way interaction was found (diagnosis x gene x subregion; F(18,90.961) = 3.387, p<0.001). This indicated that aMCC subregions exhibited diagnosis-associated differences in o.d. for each gene. Differences in PV and SST expression were identified exclusively in BA24c’ (PV: F(2,89.262) = 3.094, p=0.05 and SST: F(2,79.678) = 4.715, p=0.012), where relative to CTR, PV o.d. was decreased in SZ **(Fig. 1H**: 24c’ p=0.022) while SST expression was lower in BP (**Fig. 1I**: BA24c’ p=0.003) subjects. For VIP, diagnosis was associated with expression differences in all aMCC subdivisions (BA32’: F(2, 60.790) = 4.191, p=0.020; BA24c’: F(2,57.245) = 4.251, p=0.019; BA24ab’: F(2,71.101) = 3.245, p=0.045). VIP o.d. was lower in SZ compared to CTR in all aMCC regions (**Fig. 1J**: BA32’ p=0.011; **Fig. 1K**; BA24c’ p=0.010; **Fig. 1L**: BA24ab’ p=0.015) and decreased in BP BA32’ (**Fig. 1J**: p=0.024) and BA24c’ (**Fig. 1K**: p=0.026) vs. CTR. Differences in GAD67 expression between control and diagnosis when the aMCC was broken down by subregion did not reach significance (data not shown). Anatomical localization of aMCC mRNA alterations is schematically summarized in **Fig. S2**.

### Diagnosis-specific dPCC/RSC/ESC gene expression

Fixed effects of diagnosis along with gene and/or subregion were examined in the dPCC/RSC/ESC as described for the aMCC. Diagnosis was associated with differences in dPCC/RSC/ESC gene expression (F(2,67.248) = 5.031, p=0.009) showing that gene expression was reduced in SZ (p=0.027) and BP (p=0.020) groups vs. CTR. The significant diagnosis x gene interaction (F(9,325.498) = 106.480, p<0.001) revealed that SST (F(2,104.140) = 6.767, p=0.002) and PV (F(2, 149.348) = 8.989, p<0.001) expression changed with diagnosis. Relative to CTR, SST and PV expression was decreased in SZ (SST: p=0.018, PV: p<0.001) and BP (SST: p<0.001, PV: p=0.008) subjects. The interaction between diagnosis and gene was not significant for either GAD67 or VIP expression in the dPCC/RSC/ESC (p>0.05).

Ultimately, the significant interaction of diagnosis within each combination of gene x subregion (F(45,140)=8.474, p<0.001) indicated diagnosis-specific effects on interneuron genes within dPCC/RSC/ESC subregions. Diagnosis-related differences in SST expression were found in the dPCC (BAd23c: F(2,91.098) = 3.179, p=0.046 and BAd23b: F(2,70.106) = 3.310, p=0.042) as well as in the RSC (BA30: F(2,67.366) = 3.284, p=0.044 and BA29: F(2,68.626) = 3.483, p=0.036). As in the aMCC, SST o.d. was decreased only in BP subjects in the dPCC (**Fig. 2G**: BAd23c p=0.014; **Fig. 2H**: BAd23b p=0.013) and RSC (**Fig. 2I**: BA30 p=0.014; **Fig. 2J**: BA29 p=0.016) vs. CTR. PV expression was significantly different between diagnoses, exclusively in the RSC (BA30: F(2,65.233) = 4.533, p=0.014; BA29: F(2,56.969 = 8.217, p=0.001) and ESC (BA26: F(2,56.523) = 6.704, p=0.002), where PV expression was reduced in SZ vs. both CTR and BP in region BA30 (**Fig. 2K**, vs. CTR p=0.005; vs. BP p=0.034) and vs. CTR in BA29 (**Fig. 2L**, p<0.001), and BA26 (**Fig. 2M**, p=0.001). BP subjects also had lower PV o.d. compared to CTR in BA29 (**Fig. 2L**, p=0.013) and BA26 (**Fig. 2M**, p=0.009). Differences in the expression of GAD67 and VIP between diagnoses were not found in any dPCC/RSC/ESC subregion (p>0.05, data not shown). Anatomical localization of dPCC/RSC/ESC mRNA alterations is schematically summarized in **Fig. S2**.

## Discussion

The current study was conducted to enhance the understanding of potential GABA-ergic circuit dysfunction in the cingulate gyrus in two disorders, schizophrenia, and bipolar disorder, with both common and distinct clinical features. Employing neuroanatomical specificity, we report consistent subregional reductions in interneuron-related gene expression in the anterior and posterior cingulate gyrus. Decreased GAD67 and VIP gene expression was exclusive to aMCC and shared by both SZ and BP subjects. However, within aMCC, BA24c’ displayed diagnosis-dependent changes showing differentially reduced PV and SST mRNA in SZ and BP, respectively. In the caudal cingulate gyrus, PV reductions were shared by both SZ and BP subjects in RSC whereas SST changes, like those in aMCC, were BP-specific in dPCC and RSC.

Here we discuss the potential functional implications of these local cingulate dysregulations, using as a framework the previously proposed alterations in dynamic shifting between large-scale neurocognitive networks (DMN, SAN, and CEN) in accordance with the triple network of psychopathology.

### GAD67 Expression

Our finding of reduced aMCC GAD67 expression in SZ largely agrees with and extends two previous postmortem analyses in this region. Utilizing qPCR, Hashimoto et al. (28) showed downregulated aMCC GAD67 mRNA where its expression was not significantly altered intrinsically, but rather similarly decreased when considered along with expression in DLPFC, primary motor, and visual cortices of SZ subjects. Similar assessment of SZ subjects by Ramaker et al. (29) detected both reduced aMCC GAD67 mRNA and GABA. Taken together with the current analysis, these findings agree with SZ meta-analyses of basal aMCC GABA measurements showing a general reduction (22,23). Disruption of rostral cingulate GABA-ergic mechanisms is likely not limited to aMCC as prior findings within the anatomically adjacent pregenual ACC (pACC), a DMN hub better attuned to affective responses versus goal-directed behaviors of aMCC (1,2), highlight cortical layer II density deficits of GAD67 mRNA expressing neurons in SZ subjects (33).

Our data revealing reduced aMCC GAD67 in BP is more novel. Ramaker et al. (29), despite examining an overlapping subject cohort with the current investigation, did not uncover reduced aMCC GAD67 mRNA in BP. However, in contrast to several previous aMCC imaging analyses examining BP patients collectively demonstrating non-significant change in that region (23,34), reduced GABA was reported (29) which parallels our GAD67 results. Interestingly, in pACC, dysregulation of GABA-ergic interneurons is better associated with mood. For example, BP subjects exhibit reduced layer II GAD67 mRNA, similar to SZ (33), but also significant density decreases in non-pyramidal, presumably GABA-ergic neurons, in BP and schizoaffective disorder along with consistent albeit non-significant reductions in SZ (35,36). In contrast, comparable investigation in aMCC shows a general increase in non-specific layer VI neurons amongst both BP and SZ subjects (37).

Much of what is known regarding the potential clinical implications of altered cingulate gyrus GABA-ergic systems centers around functional analyses examining the aMCC. Concomitant with glutamate, aMCC GABA encodes evaluative feedback from conflict-driven action-outcomes and that information is subsequently used for future decision-making involving alternative actions (38,39). aMCC-mediated error processing and action monitoring is neurophysiologically assessed by the error-related negativity (ERN) corresponding to local activity in response to incorrect choices and ensuing behavioral correction (40,41). Patients with SZ, psychotic bipolar (42–44) and eurythmic BP, when controlling for depressive symptoms (45), exhibit an attenuated ERN which is connected to poor real-world functioning, negative symptom severity, and compromised executive function in psychotic disorders (43,44).

While reducing aMCC activity via GABAA receptor agonists in healthy subjects is associated with diminished ERN and action monitoring (46), lower aMCC/dPCC GABA levels are correlated with worse executive function in patients with mild cognitive impairment (47). In accord, compared to controls, first-episode psychosis patients (SZ and schizoaffective disorder) display negative correlation between aMCC GABA and total score on the Repeatable Battery for the Assessment of Neuropsychological Status, particularly in the immediate memory and language categories (48). Notably, higher aMCC GABA in first-episode psychosis patients is associated with increased local activity and Stroop task reaction time (response to incongruent and congruent stimuli) (27,28) which agrees with aMCC hyperactivation during working memory tasks in SZ (49). In sum, it is plausible that reduced aMCC GAD67 gene expression in SZ and BP subjects reflects compromised GABA-ergic neurotransmission at rest as well as stimulation of an insufficient cingulate GABA-eric signaling system associated with cognitive deficits. It would therefore be reasonable to hypothesize that reduced GABA-ergic function in this region may contribute to some of the key clinical features seen in both disorders.

Dysregulation of aMCC GABA has also been linked to aberrant large-scale neurocognitive network interactions, particularly concerning the basal ganglia. Meta-analysis demonstrates a strikingly common reduction in aMCC-caudate FC across first-episode psychosis, BP, and psychotic-bipolar patients (50). Compared to controls, reduced aMCC GABA is found in SZ spectrum disorders (22,23) and predicts a more negative FC with the caudate and putamen (26), which are SAN and CEN nodes, respectively, in first-episode psychosis (19). In psychotic bipolar disorder, aMCC-caudate FC is negatively correlated with acuteness of negative symptoms, psychopathology, and Positive and Negative Syndrome Scale (50). In addition, reduced fear perception in BP subjects is paired with decreased activation of the aMCC, caudate, and putamen (51). Of clinical significance, diminished aMCC-caudate FC also serves as a biomarker for SZ spectrum disorders as evidenced by lower strength of FC associated with better response antipsychotic drug treatment (52). Together with a SZ-related loss of correlation between aMCC GABA and FC with ventral PCC (vPCC) (53), a DMN node involved in visuospatial mnemonic processes (9), aMCC GAD67 mRNA deficits in SZ and BP subjects may contribute to psychomotor deficits through aberrant corticostriatal circuits as well blunted error processing and action monitoring, thereby supporting the triple network hypothesis of psychopathology.

### Parvalbumin Expression

Functional relevance of decreased PV gene expression in aMCC BA24c’ of SZ subjects and RSC/ectosplenial cortex (ESC) BA30/29/26 in SZ and BP can be posited from the neurophysiological properties of PV cells along with neuroanatomical and functional connectivity. Most PV interneurons are fast-spiking basket cells which utilize perisomal synapses for feedforward and feedback inhibition and generation of gamma band oscillations for synchronizing activity of pyramidal cell assemblies (27,54). Convincing arguments have been made for connecting reduced PV gene expression and GABA-ergic inhibition of DLPFC pyramidal cells to altered gamma oscillatory activity and cognitive deficits in SZ (54). Our results combined with other work showing a trend toward decreased aMCC PV mRNA in SZ (28) alludes to potentially similar deleterious effects of dysregulated BA24c’ gamma oscillatory activity on cognition. However, such alterations are probably not locally limited. Several SZ studies examining PV interneurons in BA24c, which corresponds to pACC, indicate reduced somal size and dendritic tree extent as well as increased axonal cartridges and layer V PV+ cell count (55–57). Nevertheless, although the latter concurs with heightened aMCC layer VI neuronal density in SZ (37), our results imply an overall hypofunction of BA24c’ PV interneurons, potentially affecting local gamma band oscillatory activity.

Functional imaging studies often show that BA24c’ activation associated with cognitive motor control overlaps with BA32’, thus acquiring the collective term dorsal aMCC (daMCC) as reviewed by Vogt et al (7). The functional capacity of daMCC is expansive with its activity implicated in divided attention, movement anticipation, and action initiation while also serving as the cingulate region engaged in the monitoring of ongoing action-outcomes (7) and producing ERN responses (58). Upper primate studies indicate BA24c’ function is mediated by reciprocal connections with DLPFC and premotor/supplementary motor cortex in concert with cingulospinal projections (59–62). DLPFC has been hypothesized to mediate resolve of daMCC-detected error responses (6,42) whereas DLPFC lesions alter ERN and associated corrective behavior (64). Given that DLPFC exhibits particularly strong FC with daMCC (67) and classifies as both a SAN node and CEN hub (19), it is tempting to speculate that decreased BA24c’ PV expression may point to desynchronization of aMCC-DLPFC gamma oscillatory activity during cognitive motor control tasks in SZ. However, the latter remains unclear.

More implicit evidence for daMCC-mediated cognitive motor control derives from electrical stimulation and blood flow analyses in humans showing its activation linked to body-directed arm movement, head/eye movement, and speech/speech arrest (65,66). Intertwined with this, daMCC interconnection with the insular cortex, another SAN hub, also facilitates emotional salience processing. (67). This is supported by evidence suggesting daMCC encodes value of action-outcomes for valence and arousal (68). In SZ, diminished daMCC activity is associated with impaired processing of affective asymmetry stimuli (69), anticipation of stimulus response (70), self-processing versus others (71), and executive function during N-back, go-no-go, and Stroop tasks (72). Taken together, we speculate downregulation of BA24c’ PV mRNA in SZ could be related to affective and cognitive motor control deficits. If so, aberrant daMCC communications with SAN/CEN circuits involving the insular cortex and DLPFC likely play a key role.

Within the caudal cingulate gyrus, RSC/ESC displayed the highest PV gene expression in controls while exhibiting significant BA30/29/26 reductions in SZ and BP. Importantly, interpretating these results within context of previous RSC analyses is made challenging due to potential confounds regarding various past neuroanatomical atlases/schematics incorrectly placing its location caudal to the corpus callosum splenium on the surface of the caudomedial lobule, which is actually vPCC as discussed by Vogt et al. (73). Our in situ hybridization assessment was conducted dorsal to the splenium within what we refer to as anterior RSC (aRSC) (5,73). Careful attention to brain coordinates reveals aRSC shares functional similarities with its caudal counterpart where lesions induce impairment of visuospatial memory, impaired navigation, and episodic memory (10). Indeed, aRSC activation couples with location/action recollection (9,74), anticipatory allocation of spatial attention (75), autobiographical memory (17), and memory encoding (76). In contrast, while caudal RSC is an integrated node of the self-referential DMN known to be dysregulated in SZ and BP, the role of aRSC is less clear.

Our data suggests stronger PV mRNA reductions in SZ as mean expression values were consistently lower versus BP subjects and reaching significance in RSC BA30. These finding are conceptually in-line with overlapping neurocognitive deficits in SZ and BP where SZ patients are generally more affected. Compared to controls, SZ patients demonstrate increased aRSC activity during self-reflection (77), which is influenced by visuospatial attention (78), as well as enhanced FC with the lingual gyrus (79), suggesting heightened interaction with visual systems. Along these same lines, meta-analysis of patient groups, including SZ, demonstrates increased lingual gyrus activation when evaluated while experiencing visual hallucinations (80). In addition, SZ patients with persistent auditory/verbal hallucinations exhibit increased aRSC-DMN FC and decreased aRSC-hippocampal formation FC (81,82). Reminiscent of a dysfunctional DMN, less aRSC deactivation is observed during word generation in both SZ and BP patients (83). Collectively, lower aRSC PV gene expression points to the possibility of altered gamma oscillations contributing to positive and negative symptoms, particularly in SZ.

Positioned lateral to aRSC BA29 and dorsal to the hippocampal rudiment, also referred to as indusium griseum (73), downregulation of ESC BA26 PV gene expression in SZ and BP is more difficult to interpret. Precise ESC function is unknown due, in large part, to its diminutive size being less than ideal for functional imaging assessments (73). To our knowledge, the current study is the first quantitative gene expression effort in this region. Alongside uniform aRSC PV dysregulation in BA29/30, we hypothesize ESC in SZ and BP subjects may exhibit abnormal filtering of relatively more unrefined visuospatial memory-related information. The latter is likely to be communicated to a dysfunctional aRSC in SZ and BP for further processing via ESC-BA29 interconnections.

### Somatostatin Expression

Contrasting with SZ-associated PV mRNA decreases, cingulate gyrus SST gene expression changes were strongest in BP subjects. These findings agree with comparable SZ and BP studies in orbital frontal cortex and DLPFC showing, although non-significant, lower mean levels of SST mRNA in BP versus SZ (84). In the present report, reduced SST expression was apparent in BP aMCC BA24c’, analogous to PV mRNA alterations in SZ, as well as dPCC BA23b/c where it represents the only transcriptional alteration for either diagnosis.

The differential synaptic patterns of SST+ interneuron upon pyramidal cells compared to PV+ cells dictate BA24c’ information processing likely differs by diagnosis, creating speculation and potential insight to the less sever, but overlapping psychomotor impairments observed in BP versus SZ. SST+ cells are largely Martinotti type and synapse upon distal dendrites of pyramidal cells (29) which receive afferent input from thalamic, dopaminergic, and long-range cortical afferents (85). Since SST mRNA transcription is activity dependent (86), the altered ERN amplitude in response to action monitoring displayed by BP patients (42,45) may signify reduced ability of BA24c’ SST/GABA to sharpen input from long-range cortical and subcortical inputs. SST+ neurons are important for synchronizing distant brain regions for optimal long-range cortical transmissions (29,87). More specifically, while SST interneurons have been linked to generation of theta oscillations (87,89), animal models demonstrate that disruption of SST+ cells also minimizes gamma oscillatory power in response to pharmacological and visual stimuli (89,90). Moreover, suppressing SST or PV gene transcription in medial prefrontal cortical (mPFC) produces similar SZ-like changes in social interaction and cognitive flexibility (91). Thus, similar to dysregulated BA24c’ PV in SZ, potential alterations in gamma and/or theta band oscillatory activity due to SST+ cell dysfunction may disrupt local circuits and large-scale cognitive networks mediating error/action monitoring and salience detection.

Regarding reduced SST gene expression in BA23b/c, the more dorsal aspects of dPCC, several meta-analyses associate dPCC with recognition memory and decision-making (9,92,93). As discussed by Foster et al. (2023), as opposed to decision-making contributions based upon implicit value, dPCC more likely assesses environmental prospects by integrating prospective rewards with memory-based context (9). During a cognitively demanding task, the aMCC-anchored SAN suppresses dPCC/vPCC (DMN hubs) and directs switching to the CEN for reallocation of attentional resources (12). However, dPCC BA23b/c functionally differs versus more ventral aspects. When shifting from internally directed attention to conditions of broad-external focus, although still experiencing discernable deactivation, BA23b/c continues to exhibit appreciable activity and increases FC with the CEN (8). Not until a narrower external attentional focus is required that greater deactivation ensues and FC with the CEN and DMN decreases and increases, respectively (8).

In eurythmic cognitively impaired BP patients, increased hyperconnectivity within the DMN is associated with less BA23b/c anticorrelated FC to the CEN (94), suggesting disrupted large-scale neurocognitive network dynamics. In support, lower variability, or more rigidity, of BA23b/c FC to the pACC hub of the DMN is associated with slower psychomotor speed and decreased cognitive set shifting in BP (95). Furthermore, like aMCC BA24c’ in BP, FC between dPCC and caudate is reduced (96). Altogether, our data indicate compromised BA23b/c SST+ interneurons may be connected to impairment of both motor control and allocation of attention resources needed for optimal decision-making in BP.

### Vasoactive Intestinal Peptide Expression

Unlike all other transcripts examined, reduction of VIP gene expression was observed across all quantified aMCC subregions in SZ as well as daMCC (BA32’/24c’) in BP. As a result, as determined by number of genes analyzed, the BA24c’ portion of daMCC represents the region most affected within the current aMCC analysis. In addition, decreased VIP mRNA constitutes the only altered transcript in BA32’ and BA24ab’ irrespective of diagnosis. These findings in the cingulate gyrus are novel. There are relatively few cortical studies investigating VIP expression in psychiatric illness, but two have reported reduced transcription in DLPFC and orbital frontal cortex of both SZ and BP (28) as well as the primary visual cortex node of the visuospatial working memory network in SZ subjects (97).

VIP+ GABA-ergic interneurons receive long-rang cortical afferent input and specialize in disinhibitory processes by primarily innervating dendrite-targeting SST+ cells thereby increasing the gain of excitatory pyramidal neurons (29). Elevating gain in BA24ab’ may be of particular importance regarding avoidance/escape behaviors driven by daMCC (7) considering BA24ab’ receives more appreciable input from the amydgala versus daMCC (59) and is uniquely activated by fear (7). Rodent studies show that deactivation of VIP+ interneurons in ACC or mPFC is associated with more time spent in open arms during elevated maze testing (98,99). However, upon further investigation in ACC, activity of VIP+ subpopulations preferentially increase, decrease, or remain neutral according to location in open or closed arms (99). Interestingly, the same environment-specific VIP+ cells do not stay stable over repeated trials as other specific subpopulations are replaced in later testing sessions (99).

In Humans, neuroimaging assessments reveal decreased BA24ab’ activity in SZ versus control subjects at rest (100) while also exhibiting reduced deactivation during a facial emotion discrimination requiring matching/labeling of fear and anger depictions (101). Similar studies show patients diagnosed as having high negative schizotypy with anhedonia demonstrate reduced FC between BA24ab’ and amygdala under fear and happy conditions (102). Furthermore, diminished FC between BA24ab’ and putamen is positively and negatively associated with delusion score and antipsychotic treatment, respectively (103). Thus, decreased BA24ab’ VIP expression in SZ subjects revealed here may point to the dysregulation of pyramidal cell gain control required for current and future avoidance behaviors related to fear in SZ patients.

## Conclusion

In summary, the present study uncovered divergent and overlapping reductions in interneuron subtype-specific gene expression within the rostral and caudal cingulate gyrus of SZ and BP subjects. Similar aMCC mRNA downregulations, both by gene and subregion, raise the possibility of comparable information processing impairments which may lend to altered error/action monitoring and psychomotor behavior. Caudally, patterns of gene expression changes in SZ and BP are still overlapping, yet more segregated in comparison to the aMCC. This may potentially reflect disparate circuits affecting visuospatial memory processing and attention set shifting. These results shed further light on diagnosis-related alterations in the cingulate gyrus of SZ and BP subjects.

## Acknowledgements

We thank Ms. Jennifer Fitzpatrick for technical assistance on the project. This work was supported by the Pritzker Neuropsychiatric Disorders Research Consortium.

---

**Table S1:**
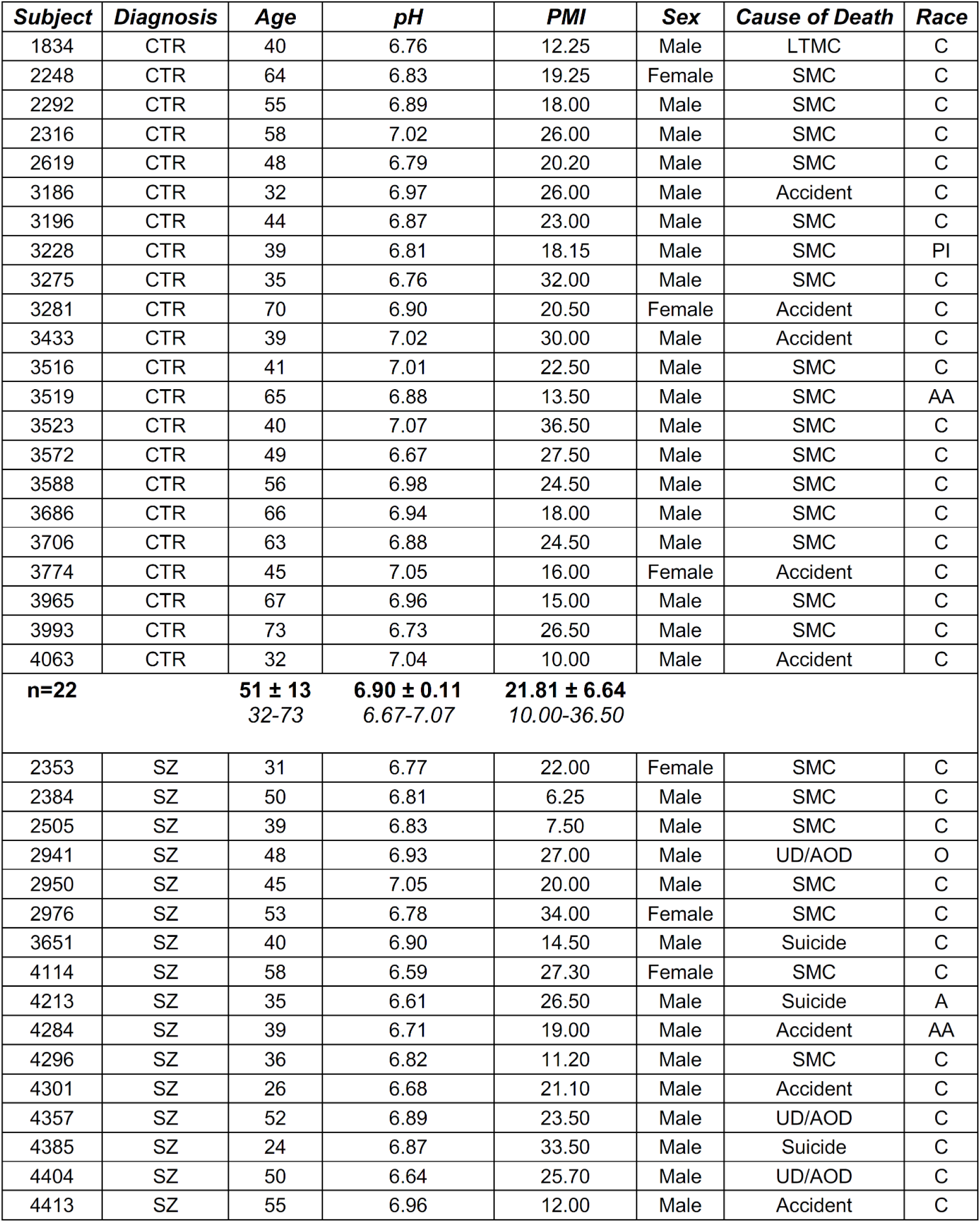
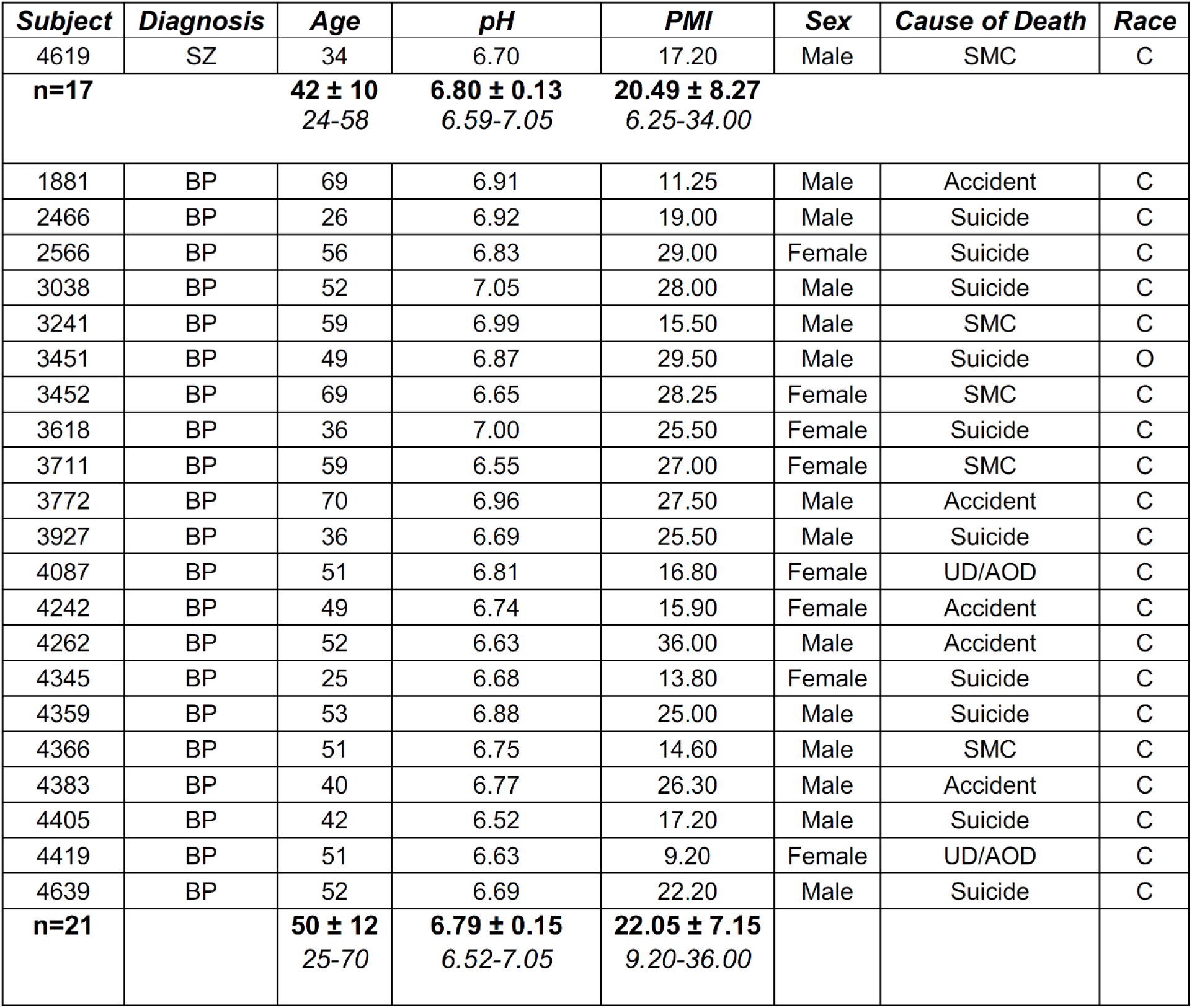
Demographic data for individual subjects studied in aMCC analyses. Subjects used in these analyses are identified by a coded subject number and grouped according to diagnosis. Relevant demographic variables including sex, age, cause of death, brain pH, post-mortem interval (PMI), and race are listed. The total number of subjects per diagnostic group is tallied at the bottom of each section in bold type. Summary information for age, pH and PMI is given at the end of each section in bold type (mean ± standard deviation), along with the range of values provided below in italics (also, see Table 1 and **Fig. S1**). aMCC: anterior midcingulate cortex; CTR: non-psychiatric control; SZ: schizophrenia; BP: bipolar disorder; SMC: sudden medical condition; UD/AOD: undetermined/accidental overdose; LTMC: long-term medical condition; PMI: postmortem interval; AA: African American; C: Caucasian; PI: Pacific Islander; A: Asian; O: Other

**Table S2:**
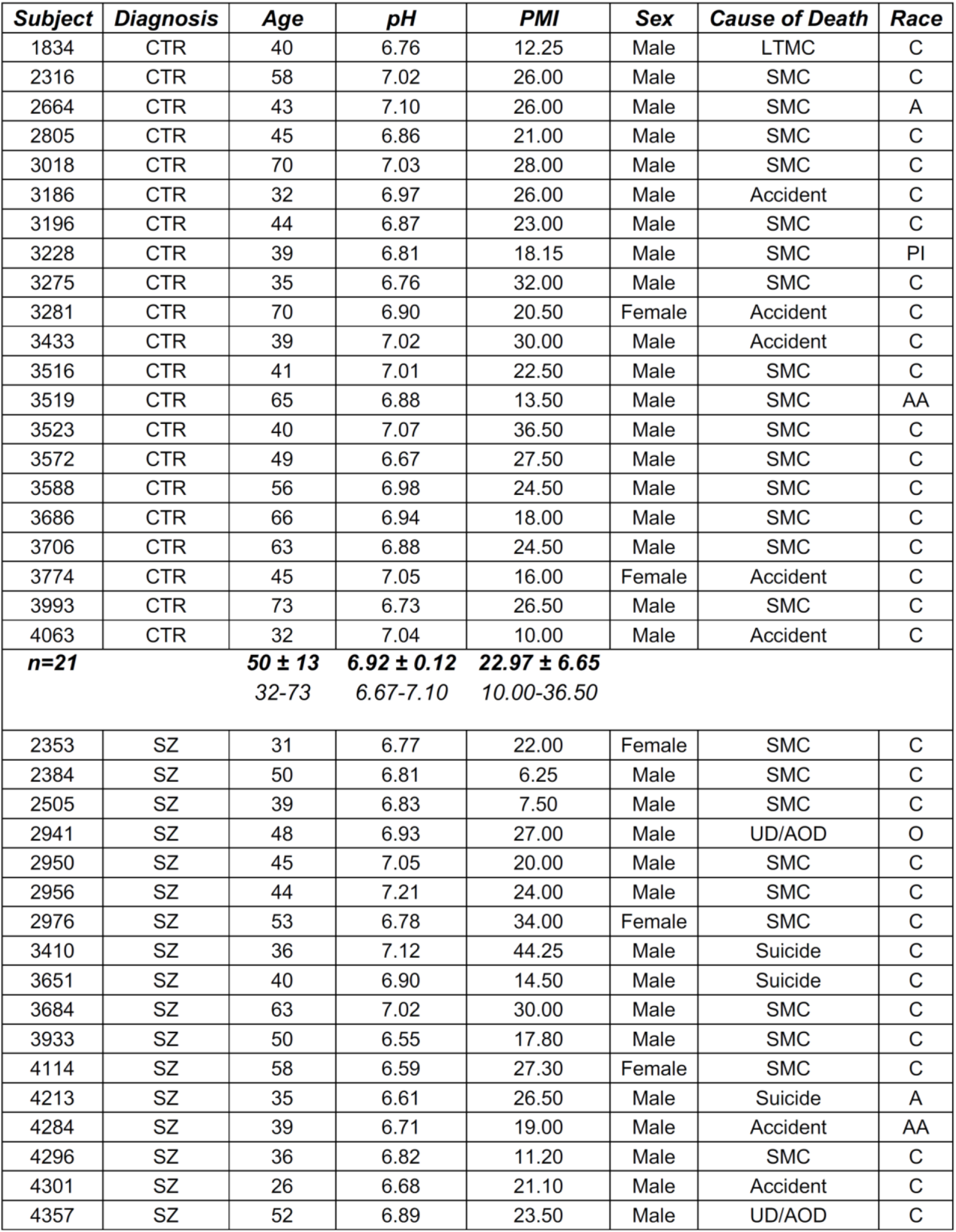
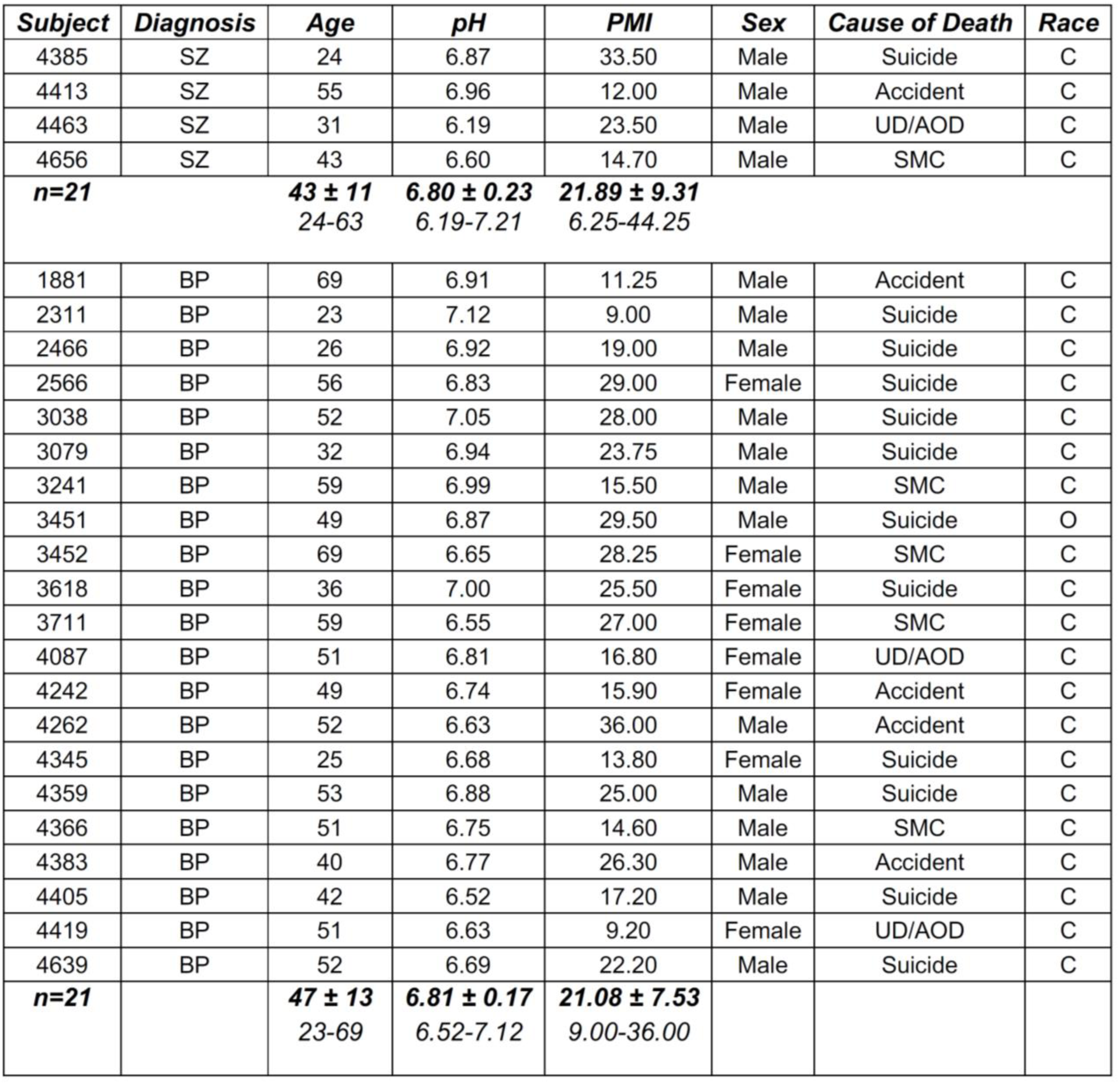
Demographic data for individual subjects studied in the dPCC/RSC/ESC analyses. This table identifies each subject by a coded subject number and associated demographics. Subjects are grouped according to diagnosis and relevant demographic variables of interest including sex, age, cause of death, brain pH, post-mortem interval (PMI), and race are indicated. The total number of subjects per diagnostic group is tallied at the bottom of each section. Summary information for CTR, SZ, and BP subject groups are listed at the bottom of each section where relevant (bolded type; mean ± standard deviation), with the range of values listed below in italics (also, see Table 1 and **Fig. S1**). dPCC: dorsal posterior cingulate cortex; RSC: retrosplenial cortex; ESC: ectosplenial cortex; CTR: non-psychiatric control; SZ: schizophrenia; BP: bipolar disorder; SMC: sudden medical condition; UD/AOD: undetermined/accidental overdose; LTMC: long-term medical condition; PMI: postmortem interval; AA: African American; C: Caucasian; PI: Pacific Islander; A: Asian; O: Other

**Table S3.**
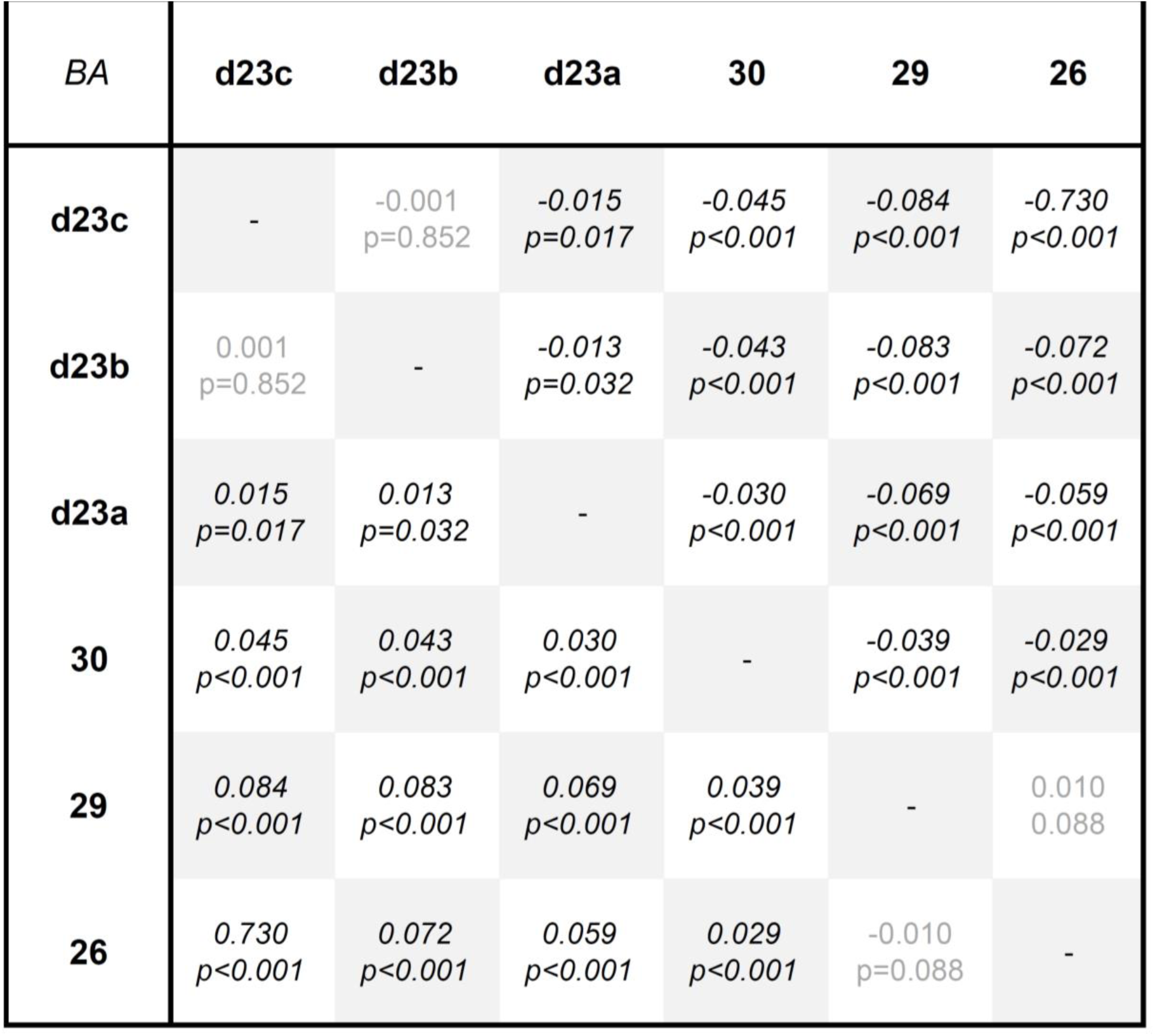
PV expression differences between dPCC/RSC/ESC subregions. The mean difference and associated p-value is indicated for each relationship between dPCC/ISH/ESC regions listed vertically compared to those along the horizontal axis. When comparing the subregion read vertically to the subregion read horizontally, a positive mean difference value (i.e., top number) means that PV expression in the vertical subregion is *greater* than expression measured in the subregion listed horizontally at the p-value indicated. Likewise, if the mean difference value is negative, it means that PV expression in the subregion read vertically is *lower* than PV expression in the subregion read horizontally at the significance level indicated. Significant differences between the vertical and horizontal subregions are italicized.

**Figure S1.**
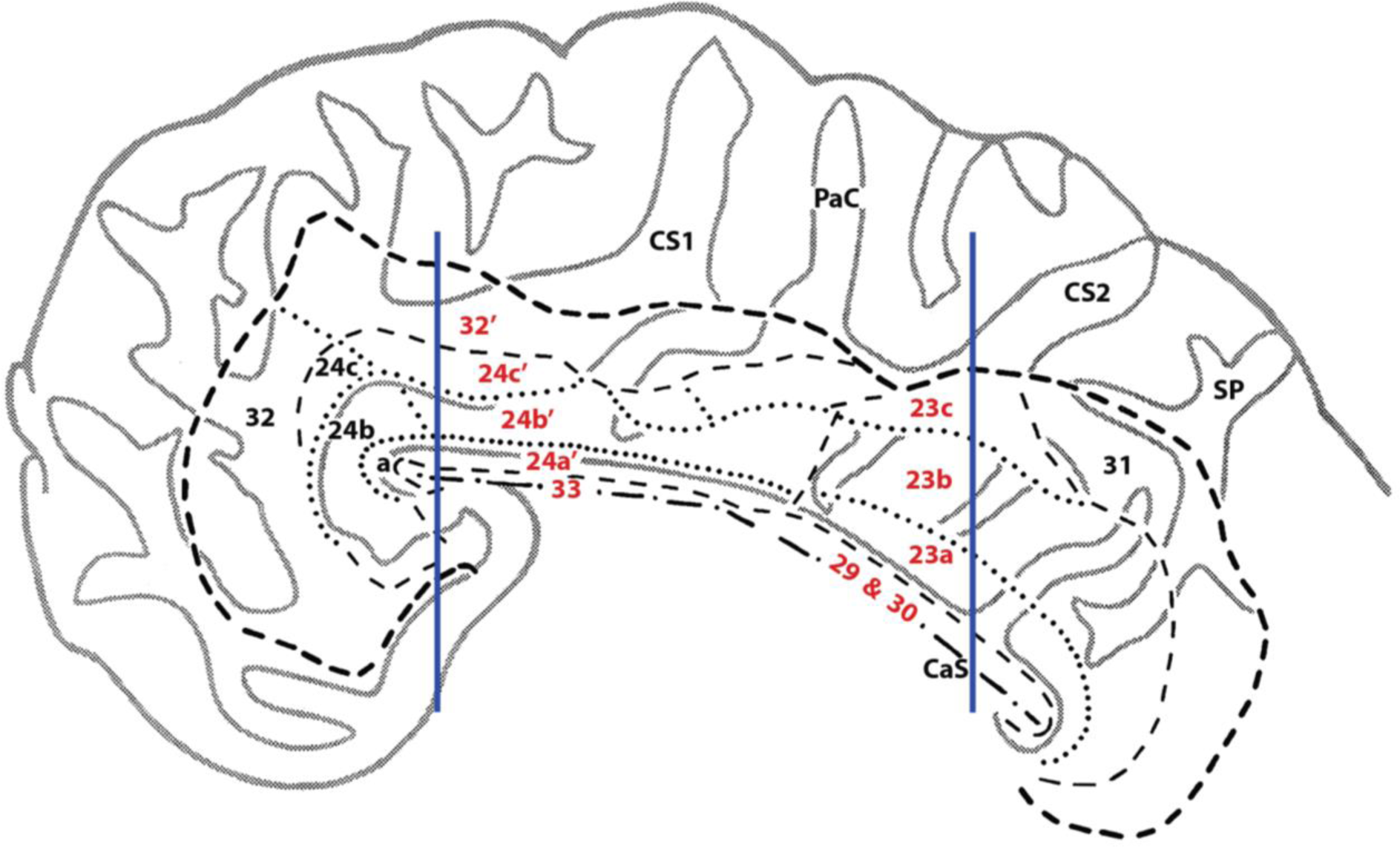
Sagittal representation of gross dissection sites used in ISH analyses. Demarcation of cingulate gyrus subdivisions in relation to gross dissection sites. Sagittal flat map depicts whole cingulate gyrus boundaries using largest bold dashed lines, major divisions outlined with thinner lines, and intra-BA subdivisions delineated by dotted lines. BAs of examined aMCC and dPCC/RSC/ESC regions are numbered in red and vertical blue lines represent dissection levels. CaS, callosal sulcus; CS, cingulate sulcus; PaC, paracentral sulcus; Sp, splenial sulcus. Modified with permission; John Wiley and Sons, J. Comp. Neurol. 359, 490–506 (1995).

**Figure S2.**
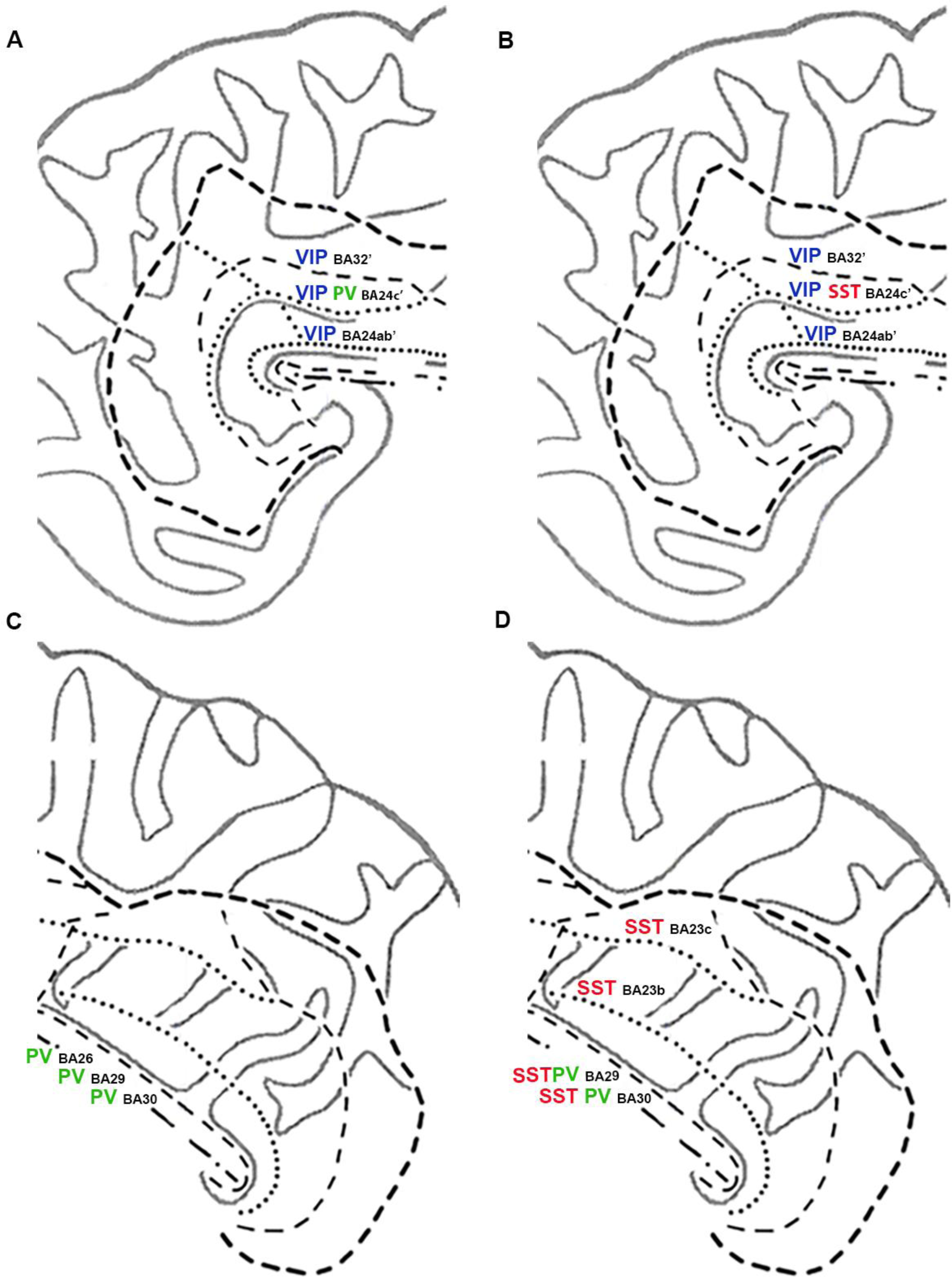
Sagittal representation showing subregional gene expression reductions. Relationship between gene expression alterations and cingulate gyrus subdivisions. All genes in BA locations represent downregulated expression as there were no increases observed the current analyses. SZ and BP gene expression decreases in aMCC are depicted in **A** and **B**, respectively, while those for the caudal cingulate gyrus are shown in **C** and **D**, respectively. Note the diagnosis-specific SST mRNA decreases in BP and the aMCC-specific alterations in GAD67/VIP within both SZ and BP. BA, Brodmann area; SST, somatostatin; PV, parvalbumin; VIP, vasoactive intestinal peptide. Modified with permission; John Wiley and Sons, J. Comp. Neurol. 359, 490–506 (1995).

**Figure S3.**
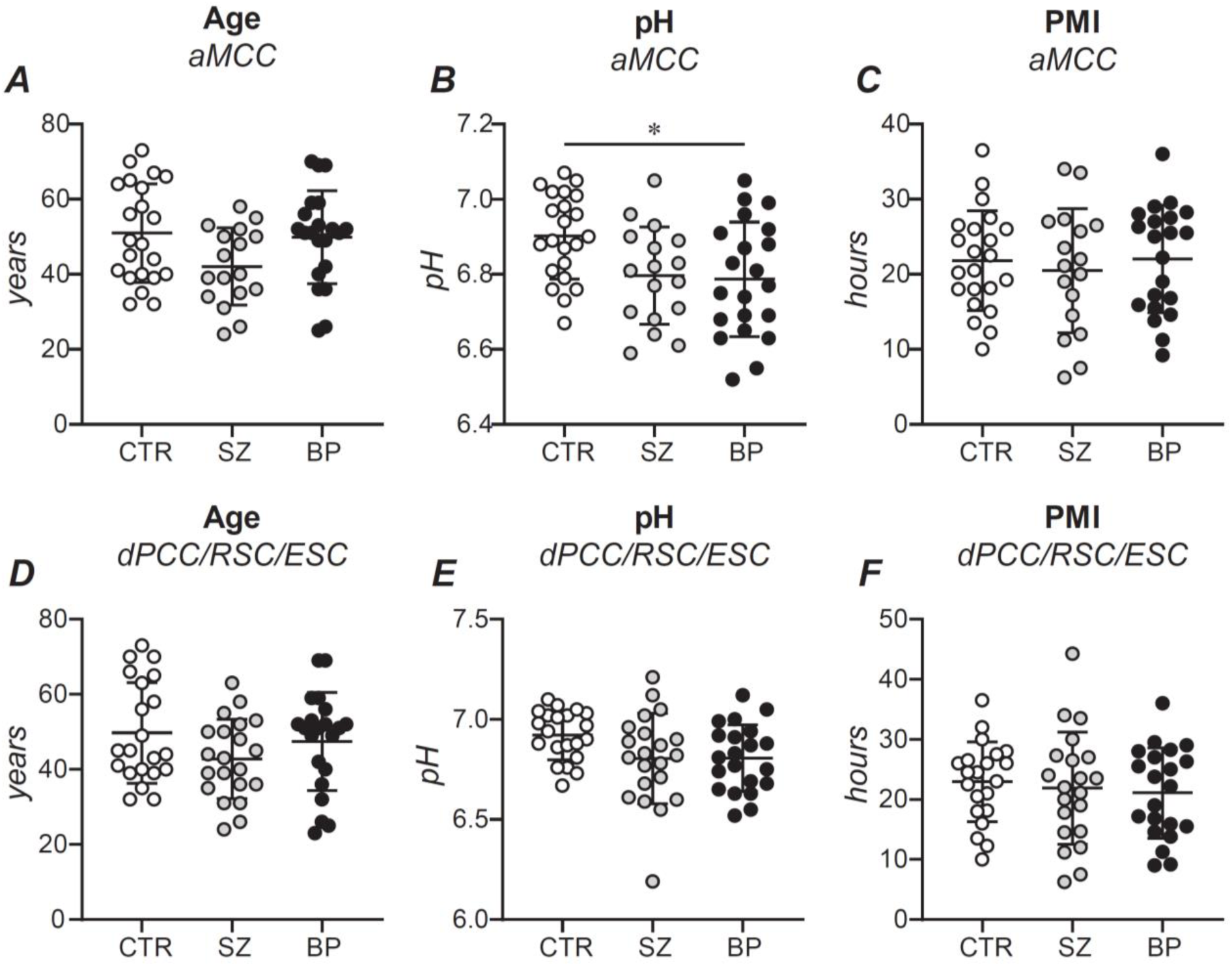
Comparison of age, pH, and PMI between diagnoses in the aMCC and dPCC/RSC/ESC. The mean differences in **A,D**) age, **B,E**) pH and **C,F**) PMI were compared between diagnoses separately in the **A-C**) aMCC and **E-F**) dPCC/RSC/ESC. Only pH (**B**) was found to differ significantly between diagnoses, with the BP subjects used in the aMCC analyses having a collectively lower average pH compared to the CTR subject group. Each circle represents a single subject; data is expressed as the mean per group and error bars represent +/-1 standard deviation. *p<0.05

**Figure S4.**
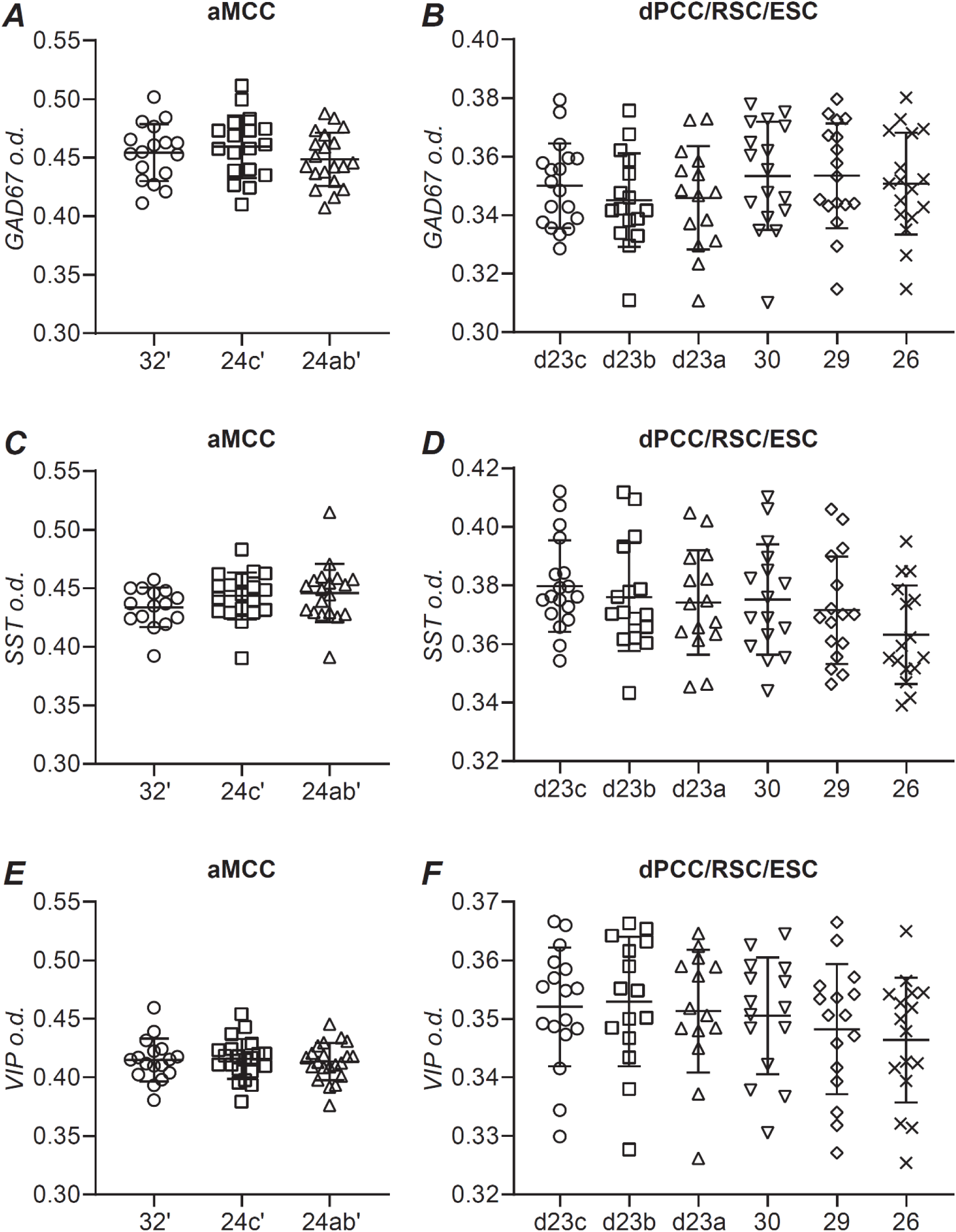
Interneuron mRNA expression in the aMCC and dPCC/RSC/ESC of control subjects. Average gene expression levels were examined within each **A,C,E)** aMCC and **B,C,F**) dPCC/RSC/ESC subregion for **A-B**) GAD67, **C-D**) SST, and **E-F**) VIP. No significant differences between subregions were detected for these genes in the aMCC or dPCC/RSC/ESC of control subjects.

## Supplementary Methods

### Neuroanatomical delineation

Cingulate lobe Brodmann’s areas (BA) and corresponding subdivisions were determined using previous publications with anatomical boundary designations (1–3). In all cases, areas were defined by gyral shape of the cortex and its relationship to anatomical landmarks (e.g., corpus callosum, cingulate sulcus). We also used gene expression patterns, especially that of PV to further define specific location (4). The rostral aMCC was localized near its border with the pACC above or just posterior to the genu of the corpus callosum. Considering histological boundaries, our gross dissection would potentially place ventral aMCC BA24a’/BA24b’ nearly adjacent to the pACC in some samples. Therefore, due to possible variance, some degree of pACC inclusion must be considered regarding quantitative analysis. To denote this, we refer to the current sampling of ventral aMCC as BA24ab’. Posterior cingulate subregions were able to be visually separated by identification of the laterally positioned RSC/ESC, indusium gresium, and spatial shift in parvalbumin gradients.

### In situ hybridization (ISH)

The processing of aMCC and PCC/RSC/ESC sections for ^35^S-labeled ISH was performed as two independent studies with each experiment utilizing hybridization of [^35^S]UTP and [^35^S]ATP-labeled cRNA riboprobes on slides taken at 500 µm intervals from the dissected blocks. Probe target, size, and accession number are as follows: GAD67, 227 nucleotides (nt), 738-964 nt of NM_000817; parvalbumin (PV), 400 nt, 60-500 nt of NM_002854; somatostatin (SST), 250 nt, 301-550 nt of NM_001048; vasoactive intestinal peptide (VIP), 624 nt, 731-1355 nt of NM_001048. Sections were hybridized and exposed to Kodak Biomax MR film (Eastman Kodak, Rochester, NY, USA). Slides from control, SZ, and BP subjects were counterbalanced within film cassettes for all experiments. The exposure time for each cRNA riboprobe was as follows: GAD67: 6 weeks, PV: 3 weeks, SST: 3 weeks, VIP: 5 weeks.

### Generation of autoradiograms

Grayscale images of the film derived ISH autoradiograms were digitized at 1600 pixels/inch using a flatbed Microtek ScanMaker 9800XL (Microtek, Carson, CA USA) and SilverFast Ai Imaging software (LaserSoft Imaging, Sarasota, FL USA). Scanned .tiff files were uploaded into Adobe Photoshop CC (2018), where adjacent groups of images within a subject (e.g., sections 1-4, sections 51-54, and sections 101-154) were overlaid and aligned in a single multi-layered .psd file prior to export as individual .tiff files for import and analysis using ImageJ (v1.50a10; National Institutes of Health, Bethesda, MD USA). Once a group of aligned files was imported into ImageJ, a “template” file was created so that comparable regions of each section were quantified for each subregion across adjacent sections.

### ISH quantification

Following ISH methods, specified areas were outlined and optical density (o.d.) of hybridization signal was obtained. It should be noted that each region of interest should be considered “enriched” BAs, since our method of ISH does not provide high resolution cellular detail. Subjects were excluded from the analyses in cases where cortical orientation could not be determined to achieve precise cortical subregion classification for at least one BA in a particular subject, within either the aMCC or dPCC/RSC/ESC. Moreover, due to potential variation in block dissection from the slab and/or cryostat artifacts (e.g., tissue tearing, folding), not all subjects contain every anatomical subregion. In any case, individual o.d. measurements of only the signal above background value were taken from 2-3 slides/subject and averaged by subregion within a subject. The o.d. of GAD67, PV, SST, and VIP mRNA is defined as follows: signal intensity above background quantified in a linear grayscale range with ImageJ generated values expressed as o.d. units.

### Pseudo-colored ISH representation

For some groups of images, a pseudo-colored composite image was created using Adobe Photoshop for visualization purposes only. In this case, the overlaid/aligned grayscale multi-layer.psd file was converted to a RGB image, but layers were not flattened. All aligned layers were then inverted, and the levels adjusted to optimize visualization of the ISH signal/noise ratio independently for each layer. The monochromatic contrast of each layer was enhanced, and a highlight color assigned to each gene (i.e., blue: GAD67, green: PV, red: SST, yellow: VIP). To visualize all ISH signals together, the blending mode was set to “lighten” for all layers excluding the bottom layer of the stack. Images were saved as multi-layer .psd and as a flattened .tiff for presentation/publication purposes.

## References

1. Rolls, E.T., 2019. The cingulate cortex and limbic systems for emotion, action, and memory. Brain Struct Funct 224, 3001–3018. 10.1007/s00429-019-01945-2

2. Vogt, B.A., 2019. The cingulate cortex in neurologic diseases: History, Structure, Overview, in: Handbook of Clinical Neurology. Elsevier, pp. 3–21. 10.1016/B978-0-444-64196-0.00001-7

3. Kuswanto, C., Chin, R., Sum, M.Y., Sengupta, S., Fagiolini, A., McIntyre, R.S., Vieta, E., Sim, K., 2016. Shared and divergent neurocognitive impairments in adult patients with schizophrenia and bipolar disorder: Whither the evidence? Neuroscience & Biobehavioral Reviews 61, 66–89. 10.1016/j.neubiorev.2015.12.002

4. Li, Y.S., Nassar, M.R., Kable, J.W., Gold, J.I., 2019. Individual Neurons in the Cingulate Cortex Encode Action Monitoring, Not Selection, during Adaptive Decision-Making. J. Neurosci. 39, 6668–6683. 10.1523/JNEUROSCI.0159-19.2019

5. Vogt, B.A., Nimchinsky, E.A., Vogt, L.J., Hof, P.R., 1995. Human cingulate cortex: Surface features, flat maps, and cytoarchitecture. J. Comp. Neurol. 359, 490–506. 10.1002/cne.903590310

6. Botvinick, M.M., 2007. Conflict monitoring and decision making: Reconciling two perspectives on anterior cingulate function. Cognitive, Affective, & Behavioral Neuroscience 7, 356–366. 10.3758/CABN.7.4.356

7. Vogt, B.A., 2016. Midcingulate cortex: Structure, connections, homologies, functions, and diseases. Journal of Chemical Neuroanatomy 74, 28–46. 10.1016/j.jchemneu.2016.01.010

8. Leech, R., Sharp, D.J., 2014. The role of the posterior cingulate cortex in cognition and disease. Brain 137, 12–32. 10.1093/brain/awt162

9. Foster, B.L., Koslov, S.R., Aponik-Gremillion, L., Monko, M.E., Hayden, B.Y., Heilbronner, S.R., 2023a. A tripartite view of the posterior cingulate cortex. Nat Rev Neurosci 24, 173–189. 10.1038/s41583-022-00661-x

10. Maguire, E., 2001. The retrosplenial contribution to human navigation: A review of lesion and neuroimaging findings. Scandinavian J Psychology 42, 225–238. 10.1111/1467-9450.00233

11. Vann, S.D., Aggleton, J.P., Maguire, E.A., 2009. What does the retrosplenial cortex do? Nat Rev Neurosci 10, 792–802. 10.1038/nrn2733

12 Menon, V., 2011. Large-scale brain networks and psychopathology: a unifying triple network model. Trends in Cognitive Sciences 15, 483–506. 10.1016/j.tics.2011.08.003

13. Touroutoglou, A., Dickerson, B.C., 2019. Cingulate-centered large-scale networks: Normal functions, aging, and neurodegenerative disease, in: Handbook of Clinical Neurology. Elsevier, pp. 113–127. 10.1016/B978-0-444-64196-0.00008-X

14. Raichle, M.E., MacLeod, A.M., Snyder, A.Z., Powers, W.J., Gusnard, D.A., Shulman, G.L., 2001. A default mode of brain function. Proc. Natl. Acad. Sci. U.S.A. 98, 676–682. 10.1073/pnas.98.2.676

15. Greicius, M.D., Krasnow, B., Reiss, A.L., Menon, V., 2003. Functional connectivity in the resting brain: A network analysis of the default mode hypothesis. Proc. Natl. Acad. Sci. U.S.A. 100, 253–258. 10.1073/pnas.0135058100

16. Fox, M.D., Snyder, A.Z., Vincent, J.L., Corbetta, M., Van Essen, D.C., Raichle, M.E., 2005. The human brain is intrinsically organized into dynamic, anticorrelated functional networks. Proc. Natl. Acad. Sci. U.S.A. 102, 9673–9678. 10.1073/pnas.0504136102

17. Spreng, R.N., Mar, R.A., Kim, A.S.N., 2009. The Common Neural Basis of Autobiographical Memory, Prospection, Navigation, Theory of Mind, and the Default Mode: A Quantitative Meta-analysis. Journal of Cognitive Neuroscience 21, 489–510. 10.1162/jocn.2008.21029

18. Buckner, R.L., DiNicola, L.M., 2019. The brain’s default network: updated anatomy, physiology, and evolving insights. Nat Rev Neurosci 20, 593–608. 10.1038/s41583-019-0212-7

19. Seeley, W.W., Menon, V., Schatzberg, A.F., Keller, J., Glover, G.H., Kenna, H., Reiss, A.L., Greicius, M.D., 2007. Dissociable Intrinsic Connectivity Networks for Salience Processing and Executive Control. J. Neurosci. 27, 2349–2356. 10.1523/JNEUROSCI.5587-06.2007

20. Nekovarova, T., Fajnerova, I., Horacek, J., Spaniel, F., 2014. Bridging disparate symptoms of schizophrenia: a triple network dysfunction theory. Front. Behav. Neurosci. 8. 10.3389/fnbeh.2014.00171

21. Yoon, S., Kim, T.D., Kim, J., Lyoo, I.K., 2021. Altered functional activity in bipolar disorder: A comprehensive review from a large-scale network perspective. Brain and Behavior 11, e01953. 10.1002/brb3.1953

22. Kumar, V., Manchegowda, S., Jacob, A., Rao, N.P., 2020. Glutamate metabolites in treatment resistant schizophrenia: A meta-analysis and systematic review of 1H-MRS studies. Psychiatry Research: Neuroimaging 300, 111080. 10.1016/j.pscychresns.2020.111080

23. Simmonite, M., Steeby, C.J., Taylor, S.F., 2023. Medial Frontal Cortex GABA Concentrations in Psychosis Spectrum and Mood Disorders: A Meta-analysis of Proton Magnetic Resonance Spectroscopy Studies. Biological Psychiatry 93, 125–136. 10.1016/j.biopsych.2022.08.004

24. Shukla, D.K., Wijtenburg, S.A., Chen, H., Chiappelli, J.J., Kochunov, P., Hong, L.E., Rowland, L.M., 2019. Anterior Cingulate Glutamate and GABA Associations on Functional Connectivity in Schizophrenia. Schizophrenia Bulletin 45, 647–658. 10.1093/schbul/sby075

25. Overbeek, G., Gawne, T.J., Reid, M.A., Salibi, N., Kraguljac, N.V., White, D.M., Lahti, A.C., 2019. Relationship Between Cortical Excitation and Inhibition and Task-Induced Activation and Deactivation: A Combined Magnetic Resonance Spectroscopy and Functional Magnetic Resonance Imaging Study at 7T in First-Episode Psychosis. Biological Psychiatry: Cognitive Neuroscience and Neuroimaging 4, 121–130. 10.1016/j.bpsc.2018.10.002

26. Overbeek, G., Gawne, T.J., Reid, M.A., Kraguljac, N.V., Lahti, A.C., 2021. A multimodal neuroimaging study investigating resting-state connectivity, glutamate, and GABA at 7 T in first-episode psychosis. jpn 46, E702–E710. 10.1503/jpn.210107

27. Tremblay, R., Lee, S., Rudy, B., 2016. GABAergic Interneurons in the Neocortex: From Cellular Properties to Circuits. Neuron 91, 260–292. 10.1016/j.neuron.2016.06.033

28. Hashimoto, T., Bazmi, H.H., Mirnics, K., Wu, Q., Sampson, A.R., Lewis, D.A., 2008. Conserved Regional Patterns of GABA-Related Transcript Expression in the Neocortex of Subjects With Schizophrenia. AJP 165, 479–489. 10.1176/appi.ajp.2007.07081223

29. Ramaker, R.C., Bowling, K.M., Lasseigne, B.N., Hagenauer, M.H., Hardigan, A.A., Davis, N.S., Gertz, J., Cartagena, P.M., Walsh, D.M., Vawter, M.P., Jones, E.G., Schatzberg, A.F., Barchas, J.D., Watson, S.J., Bunney, B.G., Akil, H., Bunney, W.E., Li, J.Z., Cooper, S.J., Myers, R.M., 2017. Post-mortem molecular profiling of three psychiatric disorders. Genome Med 9, 72. 10.1186/s13073-017-0458-5

30. Li, J.Z., Vawter, M.P., Walsh, D.M., Tomita, H., Evans, S.J., Choudary, P.V., Lopez, J.F., Avelar, A., Shokoohi, V., Chung, T., Mesarwi, O., Jones, E.G., Watson, S.J., Akil, H., Bunney, W.E., Myers, R.M., 2004. Systematic changes in gene expression in postmortem human brains associated with tissue pH and terminal medical conditions. Hum Mol Genet 13, 609–616. 10.1093/hmg/ddh065

31. Tomita, H., Vawter, M.P., Walsh, D.M., Evans, S.J., Choudary, P.V., Li, J., Overman, K.M., Atz, M.E., Myers, R.M., Jones, E.G., Watson, S.J., Akil, H., Bunney, W.E., 2004. Effect of agonal and postmortem factors on gene expression profile: quality control in microarray analyses of postmortem human brain. Biol Psychiatry 55, 346–352. 10.1016/j.biopsych.2003.10.013

32. Krolewski, D.M., Medina, A., Kerman, I.A., Bernard, R., Burke, S., Thompson, R.C., Bunney, W.E., Schatzberg, A.F., Myers, R.M., Akil, H., Jones, E.G., Watson, S.J., 2010. Expression patterns of corticotropin-releasing factor, arginine vasopressin, histidine decarboxylase, melanin-concentrating hormone, and orexin genes in the human hypothalamus. J Comp Neurol 518, 4591–4611. 10.1002/cne.22480

33. Woo, T.-U.W., Walsh, J.P., Benes, F.M., 2004b. Density of Glutamic Acid Decarboxylase 67 Messenger RNA–Containing Neurons That Express the N-Methyl-D-Aspartate Receptor Subunit NR2A in the Anterior Cingulate Cortex in Schizophrenia and Bipolar Disorder. Arch Gen Psychiatry 61, 649. 10.1001/archpsyc.61.7.649

34. Scotti-Muzzi, E., Umla-Runge, K., Soeiro-de-Souza, M.G., 2021. Anterior cingulate cortex neurometabolites in bipolar disorder are influenced by mood state and medication: A meta-analysis of 1H-MRS studies. European Neuropsychopharmacology 47, 62–73. 10.1016/j.euroneuro.2021.01.096

35. Benes, F.M., 1991. Deficits in Small Interneurons in Prefrontal and Cingulate Cortices of Schizophrenic and Schizoaffective Patients. Arch Gen Psychiatry 48, 996. 10.1001/archpsyc.1991.01810350036005

36. Benes, F.M., Vincent, S.L., Todtenkopf, M., 2001. The density of pyramidal and nonpyramidal neurons in anterior cingulate cortex of schizophrenic and bipolar subjects. Biological Psychiatry 50, 395–406. 10.1016/S0006-3223(01)01084-8

37. Chana G, Landau S, Beasley C, Everall IP, Cotter D. Two-dimensional assessment of cytoarchitecture in the anterior cingulate cortex in major depressive disorder, bipolar disorder, and schizophrenia: evidence for decreased neuronal somal size and increased neuronal density. Biological Psychiatry. 2003;53: 1086–1098. doi:10.1016/S0006-3223(03)00114-8

38. Williams ZM, Bush G, Rauch SL, Cosgrove GR, Eskandar EN. Human anterior cingulate neurons and the integration of monetary reward with motor responses. Nat Neurosci. 2004;7: 1370–1375. doi:10.1038/nn1354

39. Scholl, J., Kolling, N., Nelissen, N., Stagg, C.J., Harmer, C.J., Rushworth, M.F., 2017. Excitation and inhibition in anterior cingulate predict use of past experiences. eLife 6, e20365. 10.7554/eLife.20365

40. Gehring, W.J., Goss, B., Coles, M.G.H., Meyer, D.E., Donchin, E., 1993. A Neural System for Error Detection and Compensation. Psychol Sci 4, 385–390. 10.1111/j.1467-9280.1993.tb00586.x

41. Vanveen, V., Carter, C., 2002. The anterior cingulate as a conflict monitor: fMRI and ERP studies. Physiology & Behavior 77, 477–482. 10.1016/S0031-9384(02)00930-7

42. Minzenberg, M.J., Gomes, G.C., Yoon, J.H., Swaab, T.Y., Carter, C.S., 2014. Disrupted action monitoring in recent-onset psychosis patients with schizophrenia and bipolar disorder. Psychiatry Research: Neuroimaging 221, 114–121. 10.1016/j.pscychresns.2013.11.003

43. Martin, E.A., McCleery, A., Moore, M.M., Wynn, J.K., Green, M.F., Horan, W.P., 2018. ERP indices of performance monitoring and feedback processing in psychosis: A meta-analysis. International Journal of Psychophysiology 132, 365–378. 10.1016/j.ijpsycho.2018.08.004

44. Foti, D., Perlman, G., Bromet, E.J., Harvey, P.D., Hajcak, G., Mathalon, D.H., Kotov, R., 2021. Pathways from performance monitoring to negative symptoms and functional outcomes in psychotic disorders. Psychol. Med. 51, 2012–2022. 10.1017/S0033291720000768

45. Morsel, A.M., Morrens, M., Temmerman, A., Sabbe, B., De Bruijn, E.R., 2014. Electrophysiological (EEG) evidence for reduced performance monitoring in euthymic bipolar disorder. Bipolar Disorders 16, 820–829. 10.1111/bdi.12256

46. Riba, J., Rodríguez-Fornells, A., Münte, T.F., Barbanoj, M.J., 2005. A neurophysiological study of the detrimental effects of alprazolam on human action monitoring. Cognitive Brain Research 25, 554–565. 10.1016/j.cogbrainres.2005.08.009

47. Fu, X., Qin, M., Liu, X., Cheng, L., Zhang, L., Zhang, X., Lei, Y., Zhou, Q., Sun, P., Lin, L., Su, Y., Wang, J., 2023. Decreased GABA levels of the anterior and posterior cingulate cortex are associated with executive dysfunction in mild cognitive impairment. Front. Neurosci. 17, 1220122. 10.3389/fnins.2023.1220122

48. Reid, M.A., Forloines, M.R., Salibi, N., 2022. Reproducibility of 7-T brain spectroscopy using an ultrashort echo time STimulated Echo Acquisition Mode sequence and automated voxel repositioning. NMR in Biomedicine 35, e4631. 10.1002/nbm.4631

49. Wu, D., Jiang, T., 2020. Schizophrenia-related abnormalities in the triple network: a meta-analysis of working memory studies. Brain Imaging and Behavior 14, 971–980. 10.1007/s11682-019-00071-1

50. Liang, S., Cao, B., Deng, W., Kong, X., Zhao, L., Jin, Y., Ma, X., Wang, Y., Li, X., Wang, Q., Guo, W., Du, X., Sham, P.C., Greenshaw, A.J., Li, T., 2023. Functional dysconnectivity of anterior cingulate subregions in schizophrenia and psychotic and nonpsychotic bipolar disorder. Schizophrenia Research 254, 155–162. 10.1016/j.schres.2023.02.023

51. Killgore, W.D.S., Gruber, S.A., Yurgelun-Todd, D.A., 2008. Abnormal corticostriatal activity during fear perception in bipolar disorder. NeuroReport 19, 1523–1527. 10.1097/WNR.0b013e328310af58

52. Sarpal, D.K., Robinson, D.G., Lencz, T., Argyelan, M., Ikuta, T., Karlsgodt, K., Gallego, J.A., Kane, J.M., Szeszko, P.R., Malhotra, A.K., 2015. Antipsychotic Treatment and Functional Connectivity of the Striatum in First-Episode Schizophrenia. JAMA Psychiatry 72, 5. 10.1001/jamapsychiatry.2014.1734

53. Shukla, D.K., Wijtenburg, S.A., Chen, H., Chiappelli, J.J., Kochunov, P., Hong, L.E., Rowland, L.M., 2019. Anterior Cingulate Glutamate and GABA Associations on Functional Connectivity in Schizophrenia. Schizophrenia Bulletin 45, 647–658. 55.

54. Gonzalez-Burgos, G., Cho, R.Y., Lewis, D.A., 2015. Alterations in Cortical Network Oscillations and Parvalbumin Neurons in Schizophrenia. Biological Psychiatry 77, 1031– 1040. 10.1016/j.biopsych.2015.03.010

55. Kalus, P., Senitz, D., Beckmann, H., 1997. Altered distribution of parvalbumin-immunoreactive local circuit neurons in the anterior cingulate cortex of schizophrenic patients. Psychiatry Research: Neuroimaging 75, 49–59. 10.1016/S0925-4927(97)00020-6

56. Kalus, P., Senitz, D., Lauer, M., Beckmann, H., 1999. Inhibitory cartridge synapses in the anterior cingulate cortex of schizophrenics. Journal of Neural Transmission 106, 763– 771. 10.1007/s007020050197

57. Kalus, P., Bondzio, J., Federspiel, A., Müller, T.J., Zuschratter, W., 2002. Cell-type specific alterations of cortical interneurons in schizophrenic patients: Neuroreport 13, 713–717. 10.1097/00001756-200204160-00035

58. Van Veen, V., Cohen, J.D., Botvinick, M.M., Stenger, V.A., Carter, C.S., 2001. Anterior Cingulate Cortex, Conflict Monitoring, and Levels of Processing. NeuroImage 14, 1302– 1308. 10.1006/nimg.2001.0923

59. Vogt, B.A., Pandya, D.N., 1987. Cingulate cortex of the rhesus monkey: II. Cortical afferents. J of Comparative Neurology 262, 271–289. 10.1002/cne.902620208

60. Dum, R., Strick, P., 1991. The origin of corticospinal projections from the premotor areas in the frontal lobe. J. Neurosci. 11, 667–689. 10.1523/JNEUROSCI.11-03-00667.1991

61. Bates, J.F., Goldman-Rakic, P.S., 1993. Prefrontal connections of medial motor areas in the rhesus monkey. J of Comparative Neurology 336, 211–228. 10.1002/cne.903360205

62. Morecraft, R.J., Van Hoesen, G.W., 1993. Frontal granular cortex input to the cingulate (M3), supplementary (M2) and primary (M1) motor cortices in the rhesus monkey. J of Comparative Neurology 337, 669–689. 10.1002/cne.903370411

63. Gehring, W.J., Knight, R.T., 2000. Prefrontal–cingulate interactions in action monitoring. Nat Neurosci 3, 516–520. 10.1038/74899

64. Yu, C., Zhou, Y., Liu, Y., Jiang, T., Dong, H., Zhang, Y., Walter, M., 2011. Functional segregation of the human cingulate cortex is confirmed by functional connectivity based neuroanatomical parcellation. Neuroimage 54, 2571–2581. 10.1016/j.neuroimage.2010.11.018

65. Caruana, F., Gerbella, M., Avanzini, P., Gozzo, F., Pelliccia, V., Mai, R., Abdollahi, R.O., Cardinale, F., Sartori, I., Lo Russo, G., Rizzolatti, G., 2018. Motor and emotional behaviours elicited by electrical stimulation of the human cingulate cortex. Brain 141, 3035–3051. 10.1093/brain/awy219

66. Chassagnon, S., Minotti, L., Kremer, S., Hoffmann, D., Kahane, P., 2008. Somatosensory, motor, and reaching/grasping responses to direct electrical stimulation of the human cingulate motor areas: Clinical article. JNS 109, 593–604. 10.3171/JNS/2008/109/10/0593

67. Zhang, M., Peng, Y., 2023. Anterior insula and dorsal anterior cingulate cortex as a hub of self-regulation: combining activation likelihood estimation meta-analysis and meta-analytic connectivity modeling analysis. Brain Struct Funct 228, 1329–1345. 10.1007/s00429-023-02652-9

68. Bush KA, James GA, Privratsky AA, Fialkowski KP, Kilts CD. Action-value processing underlies the role of the dorsal anterior cingulate cortex in performance monitoring during self-regulation of affect. Shibuya K, editor. PLoS ONE. 2022;17: e0273376. doi:10.1371/journal.pone.0273376

69. Lee, S.-K., Chun, J.W., Lee, J.S., Park, H.-J., Jung, Y.-C., Seok, J.-H., Kim, J.-J., 2014. Abnormal Neural Processing during Emotional Salience Attribution of Affective Asymmetry in Patients with Schizophrenia. PLoS ONE 9, e90792. 10.1371/journal.pone.0090792

70. Quintana, J., Wong, T., Ortiz-Portillo, E., Marder, S.R., Mazziotta, J.C., 2004. Anterior cingulate dysfunction during choice anticipation in schizophrenia. Psychiatry Research: Neuroimaging 132, 117–130. 10.1016/j.pscychresns.2004.06.005

71. Potvin, S., Gamache, L., Lungu, O., 2019. A Functional Neuroimaging Meta-Analysis of Self-Related Processing in Schizophrenia. Front. Neurol. 10, 990. 10.3389/fneur.2019.00990

72. Minzenberg, M.J., Laird, A.R., Thelen, S., Carter, C.S., Glahn, D.C., 2009. Meta-analysis of 41 Functional Neuroimaging Studies of Executive Function in Schizophrenia. Arch Gen Psychiatry 66, 811. 10.1001/archgenpsychiatry.2009.91

73. Vogt, B.A., Vogt, L.J., Perl, D.P., Hof, P.R., 2001. Cytology of human caudomedial cingulate, retrosplenial, and caudal parahippocampal cortices. J Comp Neurol 438, 353–376. 10.1002/cne.1320

74. Brandt, V.C., Bergström, Z.M., Buda, M., Henson, R.N.A., Simons, J.S., 2014. Did I turn off the gas? Reality monitoring of everyday actions. Cogn Affect Behav Neurosci 14, 209–219. 10.3758/s13415-013-0189-z

75. Small, D.M., Gitelman, D.R., Gregory, M.D., Nobre, A.C., Parrish, T.B., Mesulam, M.-M., 2003. The posterior cingulate and medial prefrontal cortex mediate the anticipatory allocation of spatial attention. Neuroimage 18, 633–641. 10.1016/s1053-8119(02)00012-5

76. Kircher, T., Weis, S., Leube, D., Freymann, K., Erb, M., Jessen, F., Grodd, W., Heun, R., Krach, S., 2008. Anterior hippocampus orchestrates successful encoding and retrieval of non-relational memory: an event-related fMRI study. Eur Arch Psychiatry Clin Neurosc 258, 363–372. 10.1007/s00406-008-0805-z

77. Holt, D.J., Cassidy, B.S., Andrews-Hanna, J.R., Lee, S.M., Coombs, G., Goff, D.C., Gabrieli, J.D., Moran, J.M., 2011. An Anterior-to-Posterior Shift in Midline Cortical Activity in Schizophrenia During Self-Reflection. Biological Psychiatry 69, 415–423. 10.1016/j.biopsych.2010.10.003

78. Liu, Y., Tong, Y., Li, H., 2017. Self-reflection Orients Visual Attention Downward. Front. Psychol. 8, 1506. 10.3389/fpsyg.2017.01506

79. Wang, D., Zhou, Y., Zhuo, C., Qin, W., Zhu, J., Liu, H., Xu, L., Yu, C., 2015. Altered functional connectivity of the cingulate subregions in schizophrenia. Transl Psychiatry 5, e575–e575. 10.1038/tp.2015.69

80. Zmigrod, L., Garrison, J.R., Carr, J., Simons, J.S., 2016. The neural mechanisms of hallucinations: A quantitative meta-analysis of neuroimaging studies. Neuroscience & Biobehavioral Reviews 69, 113–123. 10.1016/j.neubiorev.2016.05.037

81. Alonso-Solís, A., Vives-Gilabert, Y., Grasa, E., Portella, M.J., Rabella, M., Sauras, R.B., Roldán, A., Núñez-Marín, F., Gómez-Ansón, B., Pérez, V., Alvarez, E., Corripio, I., 2015. Resting-state functional connectivity alterations in the default network of schizophrenia patients with persistent auditory verbal hallucinations. Schizophrenia Research 161, 261–268. 10.1016/j.schres.2014.10.047

82. Yu, H., Ying, W., Li, G., Lin, X., Jiang, D., Chen, G., Chen, S., Sun, X., Xu, Y., Ye, J., Zhuo, C., 2020. Exploring concomitant neuroimaging and genetic alterations in patients with and patients without auditory verbal hallucinations: A pilot study and mini review. J Int Med Res 48, 030006051988485. 10.1177/0300060519884856

83. Costafreda, S.G., Fu, C.H., Picchioni, M., Toulopoulou, T., McDonald, C., Kravariti, E., Walshe, M., Prata, D., Murray, R.M., McGuire, P.K., 2011. Pattern of neural responses to verbal fluency shows diagnostic specificity for schizophrenia and bipolar disorder. BMC Psychiatry 11, 18. 10.1186/1471-244X-11-18

84. Fung, S.J., Fillman, S.G., Webster, M.J., Shannon Weickert, C., 2014. Schizophrenia and bipolar disorder show both common and distinct changes in cortical interneuron markers. Schizophrenia Research 155, 26–30. 10.1016/j.schres.2014.02.021

85. Spruston, N., 2008. Pyramidal neurons: dendritic structure and synaptic integration. Nat Rev Neurosci 9, 206–221. 10.1038/nrn2286

86. Sánchez-Muñoz, I., Sánchez-Franco, F., Vallejo, M., Fernández, A., Palacios, N., Fernández, M., Cacicedo, L., 2010. Activity-dependent somatostatin gene expression is regulated by cAMP-dependent protein kinase and Ca ^2+^ -calmodulin kinase pathways. J of Neuroscience Research 88, 825–836. 10.1002/jnr.22264

87. Veit, J., Hakim, R., Jadi, M.P., Sejnowski, T.J., Adesnik, H., 2017. Cortical gamma band synchronization through somatostatin interneurons. Nat Neurosci 20, 951–959. 10.1038/nn.4562

88. Van Derveer, A.B., Bastos, G., Ferrell, A.D., Gallimore, C.G., Greene, M.L., Holmes, J.T., Kubricka, V., Ross, J.M., Hamm, J.P., 2021. A Role for Somatostatin-Positive Interneurons in Neuro-Oscillatory and Information Processing Deficits in Schizophrenia. Schizophrenia Bulletin 47, 1385–1398. 10.1093/schbul/sbaa184

89. Huang, P., Xiang, X., Chen, X., Li, H., 2020. Somatostatin Neurons Govern Theta Oscillations Induced by Salient Visual Signals. Cell Reports 33, 108415. 10.1016/j.celrep.2020.108415

90. Antonoudiou, P., Tan, Y.L., Kontou, G., Upton, A.L., Mann, E.O., 2020b. Parvalbumin and Somatostatin Interneurons Contribute to the Generation of Hippocampal Gamma Oscillations. J. Neurosci. 40, 7668–7687.

91. Perez, S.M., Boley, A., Lodge, D.J., 2019. Region specific knockdown of Parvalbumin or Somatostatin produces neuronal and behavioral deficits consistent with those observed in schizophrenia. Transl Psychiatry 9, 264. 10.1038/s41398-019-0603-6

92. Huang, Y., Hullfish, J., De Ridder, D., Vanneste, S., 2019. Meta-analysis of functional subdivisions within human posteromedial cortex. Brain Struct Funct 224, 435–452. 10.1007/s00429-018-1781-3

93. Wang, S., Tepfer, L.J., Taren, A.A., Smith, D.V., 2020. Functional parcellation of the default mode network: a large-scale meta-analysis. Sci Rep 10, 16096. 10.1038/s41598-020-72317-8

94. Fortea, L., Ysbæk-Nielsen, A.T., Macoveanu, J., Petersen, J.Z., Fisher, P.M., Kessing, L.V., Knudsen, G.M., Radua, J., Vieta, E., Miskowiak, K.W., 2023. Aberrant resting-state functional connectivity underlies cognitive and functional impairments in remitted patients with bipolar disorder. Acta Psychiatr Scand 148, 570–582. 10.1111/acps.13615

95. Nguyen, T.T., Kovacevic, S., Dev, S.I., Lu, K., Liu, T.T., Eyler, L.T., 2017. Dynamic functional connectivity in bipolar disorder is associated with executive function and processing speed: A preliminary study. Neuropsychology 31, 73–83. 10.1037/neu0000317

96. Teng, S., Lu, C.-F., Wang, P.-S., Li, C.-T., Tu, P.-C., Hung, C.-I., Su, T.-P., Wu, Y.-T., 2014. Altered Resting-State Functional Connectivity of Striatal-Thalamic Circuit in Bipolar Disorder. PLoS ONE 9, e96422. 10.1371/journal.pone.0096422 Research. 2014;155: 26–30. doi:10.1016/j.schres.2014.02.021

97. Tsubomoto, M., Kawabata, R., Zhu, X., Minabe, Y., Chen, K., Lewis, D.A., Hashimoto, T., 2019. Expression of Transcripts Selective for GABA Neuron Subpopulations across the Cortical Visuospatial Working Memory Network in the Healthy State and Schizophrenia. Cerebral Cortex 29, 3540–3550. 10.1093/cercor/bhy227

98. Lee, A.T., Cunniff, M.M., See, J.Z., Wilke, S.A., Luongo, F.J., Ellwood, I.T., Ponnavolu, S., Sohal, V.S., 2019. VIP Interneurons Contribute to Avoidance Behavior by Regulating Information Flow across Hippocampal-Prefrontal Networks. Neuron 102, 1223–1234.e4. 10.1016/j.neuron.2019.04.001

99. Johnson, C., Kretsge, L.N., Yen, W.W., Sriram, B., O’Connor, A., Liu, R.S., Jimenez, J.C., Phadke, R.A., Wingfield, K.K., Yeung, C., Jinadasa, T.J., Nguyen, T.P.H., Cho, E.S., Fuchs, E., Spevack, E.D., Velasco, B.E., Hausmann, F.S., Fournier, L.A., Brack, A., Melzer, S., Cruz-Martín, A., 2022. Highly unstable heterogeneous representations in VIP interneurons of the anterior cingulate cortex. Mol Psychiatry 27, 2602–2618. 10.1038/s41380-022-01485-y

100. Öngür, D., Lundy, M., Greenhouse, I., Shinn, A.K., Menon, V., Cohen, B.M., Renshaw, P.F., 2010. Default mode network abnormalities in bipolar disorder and schizophrenia. Psychiatry Research: Neuroimaging 183, 59–68. 10.1016/j.pscychresns.2010.04.008

101. Salgado-Pineda, P., Fakra, E., Delaveau, P., McKenna, P.J., Pomarol-Clotet, E., Blin, O., 2011. Correlated structural and functional brain abnormalities in the default mode network in schizophrenia patients. Schizophrenia Research 125, 101–109. 10.1016/j.schres.2010.10.027

102. Wang Y, Li Z, Liu W, Wei X, Jiang X, Lui SSY, et al. Negative Schizotypy and Altered Functional Connectivity During Facial Emotion Processing. Schizophrenia Bulletin. 2018;44: S491–S500. doi:10.1093/schbul/sby036

103. Raij, T.T., Mäntylä, T., Kieseppä, T., Suvisaari, J., 2015. Aberrant functioning of the putamen links delusions, antipsychotic drug dose, and compromised connectivity in first episode psychosis—Preliminary fMRI findings. Psychiatry Research: Neuroimaging 233, 201–211. 10.1016/j.pscychresns.2015.06.008

## Supplemental References

1. Vogt BA, Nimchinsky EA, Vogt LJ, et al.: Human cingulate cortex: Surface features, flat maps, and cytoarchitecture. J Comp Neurol 1995; 359:490–506

2. Vogt BA, Vogt LJ, Perl DP, et al.: Cytology of human caudomedial cingulate, retrosplenial, and caudal parahippocampal cortices. J Comp Neurol 2001; 438:353–376

3. Vogt BA, Vogt L, Laureys S: Cytology and functionally correlated circuits of human posterior cingulate areas. NeuroImage 2006; 29:452–466

4. Nimchinsky EA, Vogt BA, Morrison JH, et al.: Neurofilament and calcium-binding proteins in the human cingulate cortex. J Comp Neurol 1997; 384:597–620

