## Supplemental Tables for "Cell-Type Specific Reductions in Interneuron Gene Expression within Subregions of the Anterior and Posterior Cingulate Gyrus of Schizophrenia and Bipolar Disorder Subjects"

### Table S1: Demographic data for individual subjects studied in aMCC analyses

| *Subject* | *Diagnosis* | *Age* | *pH* | *PMI* | *Sex* | *Cause of Death* | *Race* |
|---|---|---|---|---|---|---|---|
| 1834 | CTR | 40 | 6.76 | 12.25 | Male | LTMC | C |
| 2248 | CTR | 64 | 6.83 | 19.25 | Female | SMC | C |
| 2292 | CTR | 55 | 6.89 | 18.00 | Male | SMC | C |
| 2316 | CTR | 58 | 7.02 | 26.00 | Male | SMC | C |
| 2619 | CTR | 48 | 6.79 | 20.20 | Male | SMC | C |
| 3186 | CTR | 32 | 6.97 | 26.00 | Male | Accident | C |
| 3196 | CTR | 44 | 6.87 | 23.00 | Male | SMC | C |
| 3228 | CTR | 39 | 6.81 | 18.15 | Male | SMC | PI |
| 3275 | CTR | 35 | 6.76 | 32.00 | Male | SMC | C |
| 3281 | CTR | 70 | 6.90 | 20.50 | Female | Accident | C |
| 3433 | CTR | 39 | 7.02 | 30.00 | Male | Accident | C |
| 3516 | CTR | 41 | 7.01 | 22.50 | Male | SMC | C |
| 3519 | CTR | 65 | 6.88 | 13.50 | Male | SMC | AA |
| 3523 | CTR | 40 | 7.07 | 36.50 | Male | SMC | C |
| 3572 | CTR | 49 | 6.67 | 27.50 | Male | SMC | C |
| 3588 | CTR | 56 | 6.98 | 24.50 | Male | SMC | C |
| 3686 | CTR | 66 | 6.94 | 18.00 | Male | SMC | C |
| 3706 | CTR | 63 | 6.88 | 24.50 | Male | SMC | C |
| 3774 | CTR | 45 | 7.05 | 16.00 | Female | Accident | C |
| 3965 | CTR | 67 | 6.96 | 15.00 | Male | SMC | C |
| 3993 | CTR | 73 | 6.73 | 26.50 | Male | SMC | C |
| 4063 | CTR | 32 | 7.04 | 10.00 | Male | Accident | C |
| **n=22** | | **51 ± 13**<br>*32-73* | **6.90 ± 0.11**<br>*6.67-7.07* | **21.81 ± 6.64**<br>*10.00-36.50* | | | |
| 2353 | SZ | 31 | 6.77 | 22.00 | Female | SMC | C |
| 2384 | SZ | 50 | 6.81 | 6.25 | Male | SMC | C |
| 2505 | SZ | 39 | 6.83 | 7.50 | Male | SMC | C |
| 2941 | SZ | 48 | 6.93 | 27.00 | Male | UD/AOD | O |
| 2950 | SZ | 45 | 7.05 | 20.00 | Male | SMC | C |
| 2976 | SZ | 53 | 6.78 | 34.00 | Female | SMC | C |
| 3651 | SZ | 40 | 6.90 | 14.50 | Male | Suicide | C |
| 4114 | SZ | 58 | 6.59 | 27.30 | Female | SMC | C |
| 4213 | SZ | 35 | 6.61 | 26.50 | Male | Suicide | A |
| 4284 | SZ | 39 | 6.71 | 19.00 | Male | Accident | AA |
| 4296 | SZ | 36 | 6.82 | 11.20 | Male | SMC | C |
| 4301 | SZ | 26 | 6.68 | 21.10 | Male | Accident | C |
| 4357 | SZ | 52 | 6.89 | 23.50 | Male | UD/AOD | C |
| 4385 | SZ | 24 | 6.87 | 33.50 | Male | Suicide | C |
| 4404 | SZ | 50 | 6.64 | 25.70 | Male | UD/AOD | C |
| 4413 | SZ | 55 | 6.96 | 12.00 | Male | Accident | C |

| ***Subject*** | ***Diagnosis*** | ***Age*** | ***pH*** | ***PMI*** | ***Sex*** | ***Cause of Death*** | ***Race*** |
|---|---|---|---|---|---|---|---|
| 4619 | SZ | 34 | 6.70 | 17.20 | Male | SMC | C |
| **n=17** | | **42 ± 10**<br>*24-58* | **6.80 ± 0.13**<br>*6.59-7.05* | **20.49 ± 8.27**<br>*6.25-34.00* | | | |
| 1881 | BP | 69 | 6.91 | 11.25 | Male | Accident | C |
| 2466 | BP | 26 | 6.92 | 19.00 | Male | Suicide | C |
| 2566 | BP | 56 | 6.83 | 29.00 | Female | Suicide | C |
| 3038 | BP | 52 | 7.05 | 28.00 | Male | Suicide | C |
| 3241 | BP | 59 | 6.99 | 15.50 | Male | SMC | C |
| 3451 | BP | 49 | 6.87 | 29.50 | Male | Suicide | O |
| 3452 | BP | 69 | 6.65 | 28.25 | Female | SMC | C |
| 3618 | BP | 36 | 7.00 | 25.50 | Female | Suicide | C |
| 3711 | BP | 59 | 6.55 | 27.00 | Female | SMC | C |
| 3772 | BP | 70 | 6.96 | 27.50 | Male | Accident | C |
| 3927 | BP | 36 | 6.69 | 25.50 | Male | Suicide | C |
| 4087 | BP | 51 | 6.81 | 16.80 | Female | UD/AOD | C |
| 4242 | BP | 49 | 6.74 | 15.90 | Female | Accident | C |
| 4262 | BP | 52 | 6.63 | 36.00 | Male | Accident | C |
| 4345 | BP | 25 | 6.68 | 13.80 | Female | Suicide | C |
| 4359 | BP | 53 | 6.88 | 25.00 | Male | Suicide | C |
| 4366 | BP | 51 | 6.75 | 14.60 | Male | SMC | C |
| 4383 | BP | 40 | 6.77 | 26.30 | Male | Accident | C |
| 4405 | BP | 42 | 6.52 | 17.20 | Male | Suicide | C |
| 4419 | BP | 51 | 6.63 | 9.20 | Female | UD/AOD | C |
| 4639 | BP | 52 | 6.69 | 22.20 | Male | Suicide | C |
| **n=21** | | **50 ± 12**<br>*25-70* | **6.79 ± 0.15**<br>*6.52-7.05* | **22.05 ± 7.15**<br>*9.20-36.00* | | | |

**Table S1** (continued)

***Table S2: Demographic data for subjects studied in dPCC/RSC/ESC analyses***

| *Subject* | *Diagnosis* | *Age* | *pH* | *PMI* | *Sex* | *Cause of Death* | *Race* |
|---|---|---|---|---|---|---|---|
| 1834 | CTR | 40 | 6.76 | 12.25 | Male | LTMC | C |
| 2316 | CTR | 58 | 7.02 | 26.00 | Male | SMC | C |
| 2664 | CTR | 43 | 7.10 | 26.00 | Male | SMC | A |
| 2805 | CTR | 45 | 6.86 | 21.00 | Male | SMC | C |
| 3018 | CTR | 70 | 7.03 | 28.00 | Male | SMC | C |
| 3186 | CTR | 32 | 6.97 | 26.00 | Male | Accident | C |
| 3196 | CTR | 44 | 6.87 | 23.00 | Male | SMC | C |
| 3228 | CTR | 39 | 6.81 | 18.15 | Male | SMC | PI |
| 3275 | CTR | 35 | 6.76 | 32.00 | Male | SMC | C |
| 3281 | CTR | 70 | 6.90 | 20.50 | Female | Accident | C |
| 3433 | CTR | 39 | 7.02 | 30.00 | Male | Accident | C |
| 3516 | CTR | 41 | 7.01 | 22.50 | Male | SMC | C |
| 3519 | CTR | 65 | 6.88 | 13.50 | Male | SMC | AA |
| 3523 | CTR | 40 | 7.07 | 36.50 | Male | SMC | C |
| 3572 | CTR | 49 | 6.67 | 27.50 | Male | SMC | C |
| 3588 | CTR | 56 | 6.98 | 24.50 | Male | SMC | C |
| 3686 | CTR | 66 | 6.94 | 18.00 | Male | SMC | C |
| 3706 | CTR | 63 | 6.88 | 24.50 | Male | SMC | C |
| 3774 | CTR | 45 | 7.05 | 16.00 | Female | Accident | C |
| 3993 | CTR | 73 | 6.73 | 26.50 | Male | SMC | C |
| 4063 | CTR | 32 | 7.04 | 10.00 | Male | Accident | C |
| ***n=21*** | | ***50 ± 13***<br>*32-73* | ***6.92 ± 0.12***<br>*6.67-7.10* | ***22.97 ± 6.65***<br>*10.00-36.50* | | | |
| 2353 | SZ | 31 | 6.77 | 22.00 | Female | SMC | C |
| 2384 | SZ | 50 | 6.81 | 6.25 | Male | SMC | C |
| 2505 | SZ | 39 | 6.83 | 7.50 | Male | SMC | C |
| 2941 | SZ | 48 | 6.93 | 27.00 | Male | UD/AOD | O |
| 2950 | SZ | 45 | 7.05 | 20.00 | Male | SMC | C |
| 2956 | SZ | 44 | 7.21 | 24.00 | Male | SMC | C |
| 2976 | SZ | 53 | 6.78 | 34.00 | Female | SMC | C |
| 3410 | SZ | 36 | 7.12 | 44.25 | Male | Suicide | C |
| 3651 | SZ | 40 | 6.90 | 14.50 | Male | Suicide | C |
| 3684 | SZ | 63 | 7.02 | 30.00 | Male | SMC | C |
| 3933 | SZ | 50 | 6.55 | 17.80 | Male | SMC | C |
| 4114 | SZ | 58 | 6.59 | 27.30 | Female | SMC | C |
| 4213 | SZ | 35 | 6.61 | 26.50 | Male | Suicide | A |
| 4284 | SZ | 39 | 6.71 | 19.00 | Male | Accident | AA |
| 4296 | SZ | 36 | 6.82 | 11.20 | Male | SMC | C |
| 4301 | SZ | 26 | 6.68 | 21.10 | Male | Accident | C |
| 4357 | SZ | 52 | 6.89 | 23.50 | Male | UD/AOD | C |

| ***Subject*** | ***Diagnosis*** | ***Age*** | ***pH*** | ***PMI*** | ***Sex*** | ***Cause of Death*** | ***Race*** |
|---|---|---|---|---|---|---|---|
| 4385 | SZ | 24 | 6.87 | 33.50 | Male | Suicide | C |
| 4413 | SZ | 55 | 6.96 | 12.00 | Male | Accident | C |
| 4463 | SZ | 31 | 6.19 | 23.50 | Male | UD/AOD | C |
| 4656 | SZ | 43 | 6.60 | 14.70 | Male | SMC | C |
| ***n=21*** | | ***43 ± 11***<br>*24-63* | ***6.80 ± 0.23***<br>*6.19-7.21* | ***21.89 ± 9.31***<br>*6.25-44.25* | | | |
| 1881 | BP | 69 | 6.91 | 11.25 | Male | Accident | C |
| 2311 | BP | 23 | 7.12 | 9.00 | Male | Suicide | C |
| 2466 | BP | 26 | 6.92 | 19.00 | Male | Suicide | C |
| 2566 | BP | 56 | 6.83 | 29.00 | Female | Suicide | C |
| 3038 | BP | 52 | 7.05 | 28.00 | Male | Suicide | C |
| 3079 | BP | 32 | 6.94 | 23.75 | Male | Suicide | C |
| 3241 | BP | 59 | 6.99 | 15.50 | Male | SMC | C |
| 3451 | BP | 49 | 6.87 | 29.50 | Male | Suicide | O |
| 3452 | BP | 69 | 6.65 | 28.25 | Female | SMC | C |
| 3618 | BP | 36 | 7.00 | 25.50 | Female | Suicide | C |
| 3711 | BP | 59 | 6.55 | 27.00 | Female | SMC | C |
| 4087 | BP | 51 | 6.81 | 16.80 | Female | UD/AOD | C |
| 4242 | BP | 49 | 6.74 | 15.90 | Female | Accident | C |
| 4262 | BP | 52 | 6.63 | 36.00 | Male | Accident | C |
| 4345 | BP | 25 | 6.68 | 13.80 | Female | Suicide | C |
| 4359 | BP | 53 | 6.88 | 25.00 | Male | Suicide | C |
| 4366 | BP | 51 | 6.75 | 14.60 | Male | SMC | C |
| 4383 | BP | 40 | 6.77 | 26.30 | Male | Accident | C |
| 4405 | BP | 42 | 6.52 | 17.20 | Male | Suicide | C |
| 4419 | BP | 51 | 6.63 | 9.20 | Female | UD/AOD | C |
| 4639 | BP | 52 | 6.69 | 22.20 | Male | Suicide | C |
| ***n=21*** | | ***47 ± 13***<br>*23-69* | ***6.81 ± 0.17***<br>*6.52-7.12* | ***21.08 ± 7.53***<br>*9.00-36.00* | | | |

**Table S2**

Krolewski and Waselus, et al.,

*Table S3. PV expression differences between dPCC/RSC/ESC subregions*

| *BA* | **d23c** | **d23b** | **d23a** | **30** | **29** | **26** |
|---|---|---|---|---|---|---|
| **d23c** | - | -0.001 p=0.852 | *-0.015 p=0.017* | *-0.045 p<0.001* | *-0.084 p<0.001* | *-0.730 p<0.001* |
| **d23b** | 0.001 p=0.852 | - | *-0.013 p=0.032* | *-0.043 p<0.001* | *-0.083 p<0.001* | *-0.072 p<0.001* |
| **d23a** | *0.015 p=0.017* | *0.013 p=0.032* | - | *-0.030 p<0.001* | *-0.069 p<0.001* | *-0.059 p<0.001* |
| **30** | *0.045 p<0.001* | *0.043 p<0.001* | *0.030 p<0.001* | - | *-0.039 p<0.001* | *-0.029 p<0.001* |
| **29** | *0.084 p<0.001* | *0.083 p<0.001* | *0.069 p<0.001* | *0.039 p<0.001* | - | 0.010 0.088 |
| **26** | *0.730 p<0.001* | *0.072 p<0.001* | *0.059 p<0.001* | *0.029 p<0.001* | -0.010 p=0.088 | - |

**Table S3**
Krolewski and Waselus, et al.,
