## Supplemental Figures for "Cell-Type Specific Reductions in Interneuron Gene Expression within Subregions of the Anterior and Posterior Cingulate Gyrus of Schizophrenia and Bipolar Disorder Subjects"

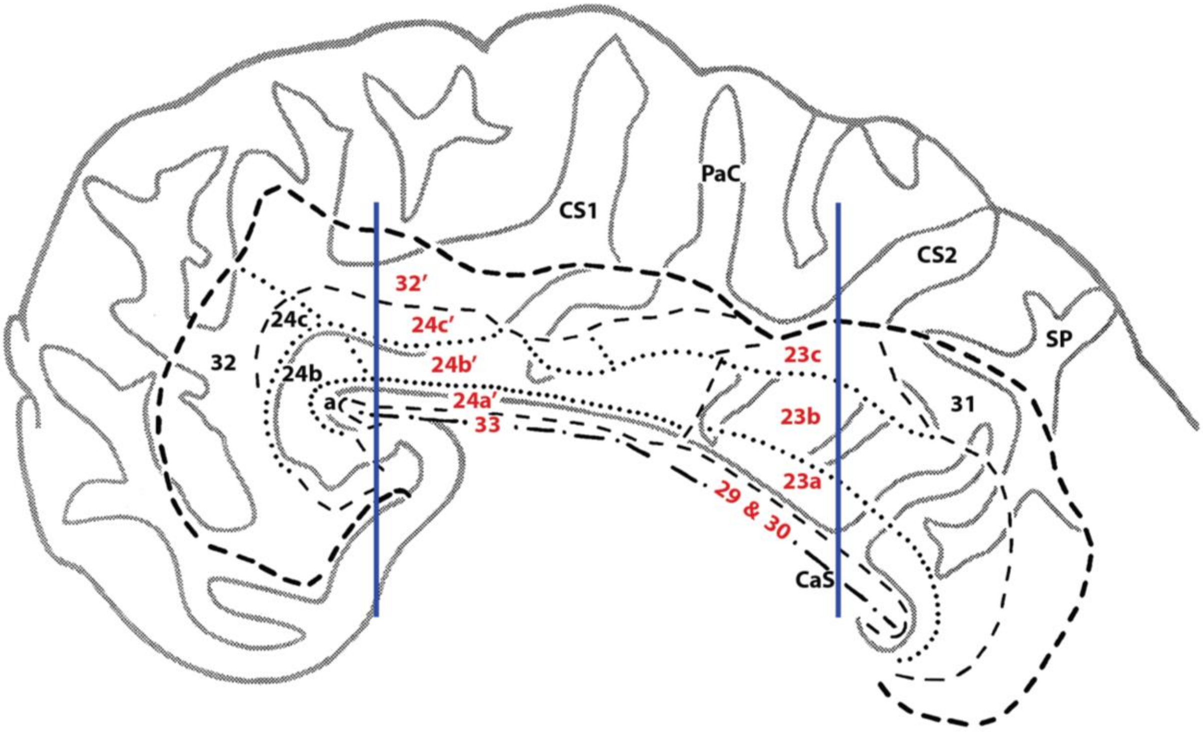


**Figure S1**
Krolewski and Waselus, et al.,

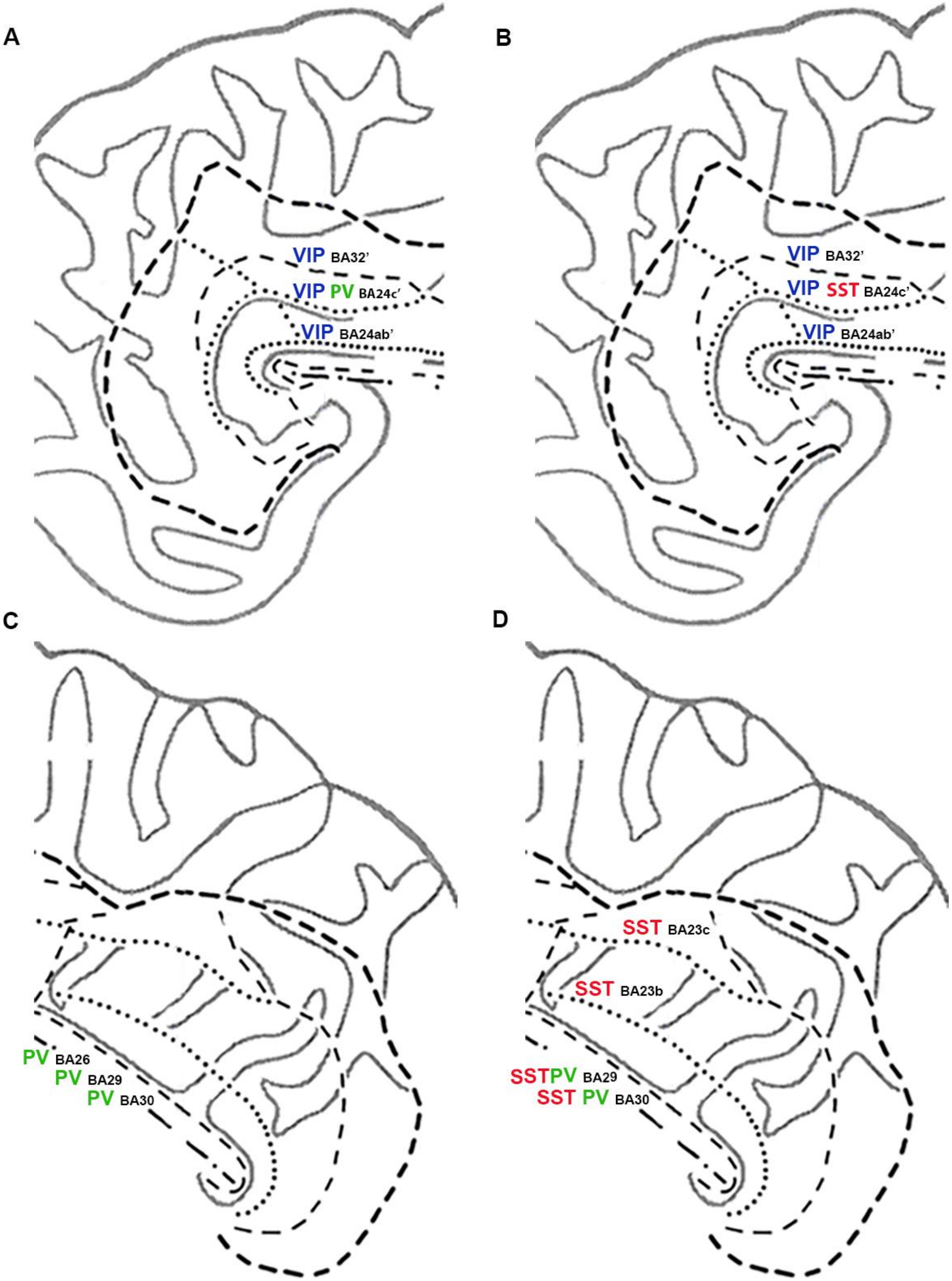


**Figure S2**

Krolewski and Waselus et. al.,

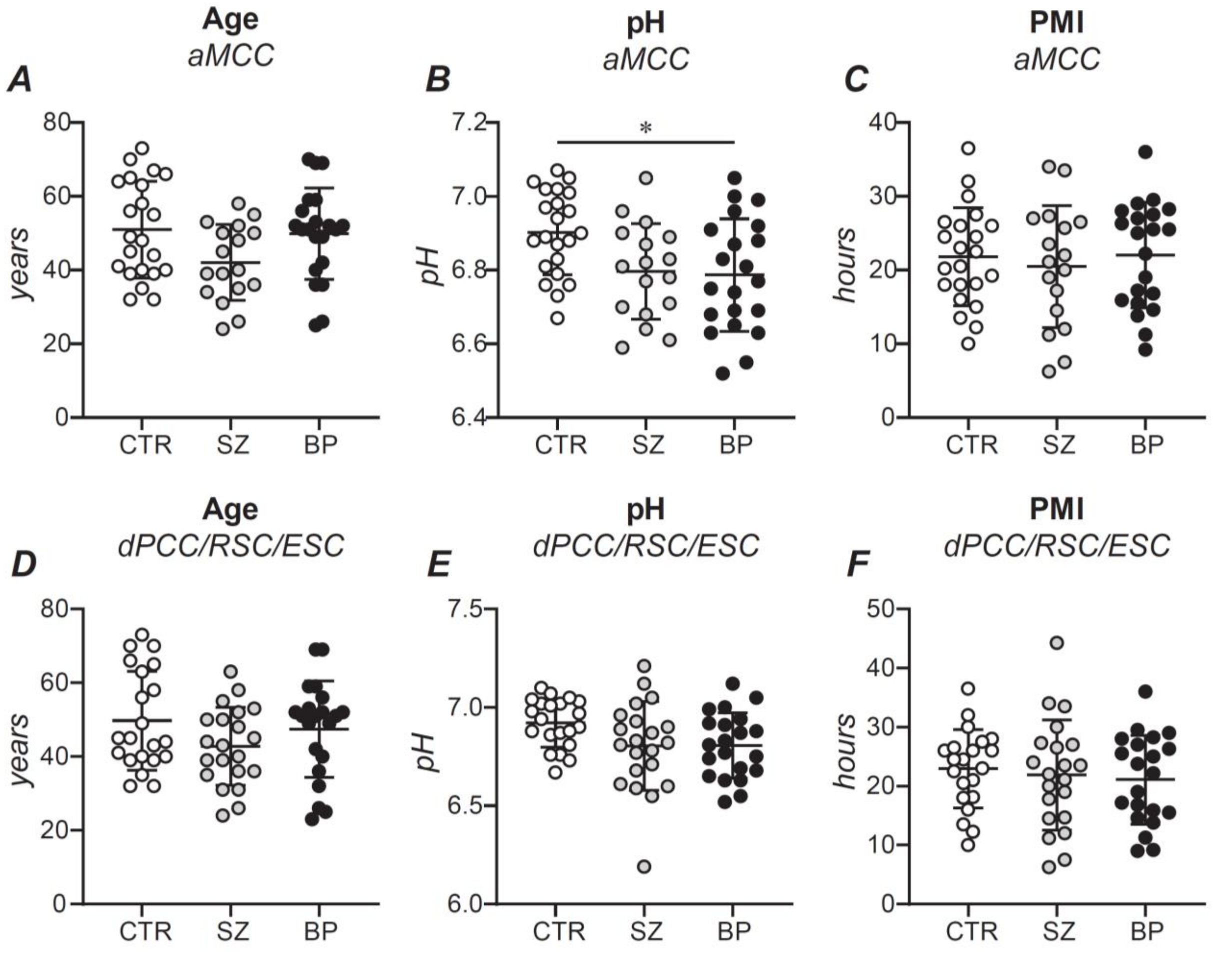


**Figure S3**

Krolewski and Waselus et. al.,

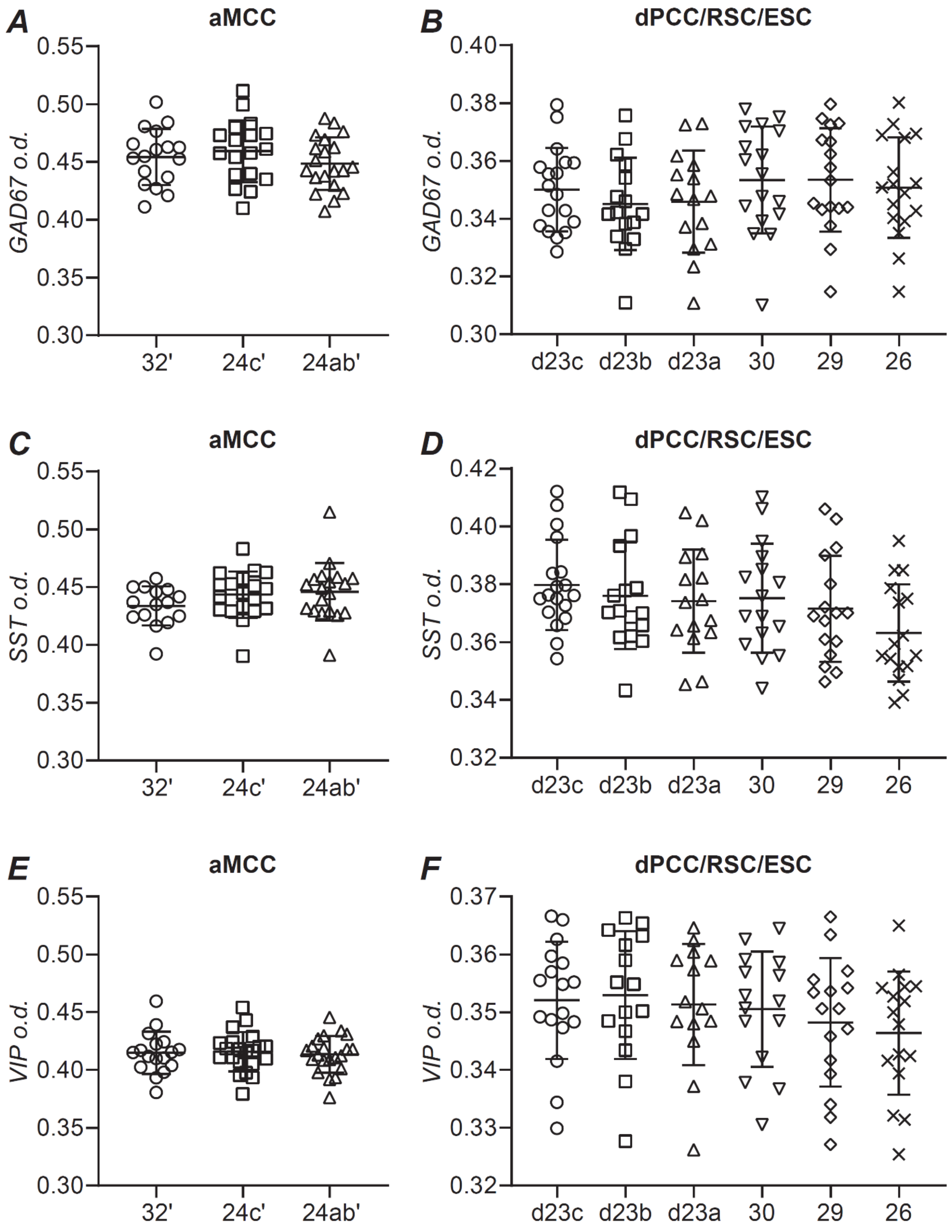


**Figure S4**

Krolewski and Waselus et. al.,
